# Two Pleckstrin homology domains scaffold SNAP-47-containing SNARE complexes

**DOI:** 10.64898/2026.10.15.759244

**Authors:** Ismar Alickovic, Molly Wise, Rain Tela, Caleb Fausnaught, Lyndsey Lee, Connor Beasley, Christopher Carr, Camilla Carr, Luke Schwindt, John Peters

**Affiliations:** Department of Biology, University of Richmond, Richmond, VA 23173

## Abstract

Long-term potentiation requires the regulated insertion of AMPA receptors into the postsynaptic membrane of neurons, a process mediated by a SNARE complex containing SNAP-47, syntaxin-3, and synaptobrevin-2. Compared to other SNAP proteins, SNAP-47 exhibits slower fusion kinetics and contains uncharacterized regions whose functions remain unclear. Here, we integrated biochemical approaches with artificial intelligence structure prediction tools and molecular dynamics to investigate the architecture of SNAP-47-containing SNARE complexes. AlphaFold3 predictions of the ternary SNAP-47-containing SNARE complex revealed one known Pleckstrin homology domain (PH1) and one previously unidentified PH domain (PH2), which we validated experimentally. BioEmu and molecular dynamics simulations indicated that PH1 and PH2 form a stable and energetically favorable interface. In this bound state, PH1 stably interacted with the SNARE coiled coil, and in silico acetylation at a lysine whose acetylation is known to impair SNAP-47 function disrupted this PH1—SNARE interface. AMPAR insertion via SNAP-47-containing SNARE complexes is triggered by the calcium sensor synaptotagmin, and we observed that PH1, PH2, and synaptotagmin-1 can simultaneously engage with the plasma membrane, consistent with a pro-fusion conformation. Together, we provide new mechanistic insight into how PH domains regulate SNAP-47-mediated membrane fusion during synaptic plasticity.

## Introduction

NMDA-receptor-dependent long-term potentiation (LTP), a cellular manifestation of memory formation, requires the delivery of AMPA receptors (AMPARs) to the surface of postsynaptic neurons^1–5^. Insertion of AMPARs is mediated by a SNARE complex containing SNAP-47, syntaxin-3 (Stx-3), and synaptobrevin-2 (Syb-2), the formation of which provides the energy required to fuse AMPAR-containing vesicles with the membrane^6–8^. SNARE complex formation is modulated by many proteins, including complexin and the calcium sensor synaptotagmin, which bind to intermediate SNARE complexes to regulate fusion^9,10^. Much of our understanding of SNARE regulation comes from work describing the assembly, triggering, and disassembly of the presynaptic SNAP-25—Syb-2—Stx-1 complex, which is involved in neurotransmitter release^11^. Initially, the complex adopts a trans conformation in which the Syb-2 transmembrane domain is embedded in the membrane of a synaptic vesicle, and the Stx-1 transmembrane domain is embedded in the plasma membrane^9^. SNAP-25 is tethered to the plasma membrane via palmitoylation of a poly-cysteine region^12^. In the presynaptic system, complexin binds to the groove formed between Stx-1 and Syb-2 within the ternary complex, where it both prevents spontaneous vesicle release and facilitates vesicle fusion^13–15^. Following complexin binding, the calcium sensor synaptotagmin-1 (Syt-1) binds to its primary binding site, which is mostly composed of residues from the two SNAP-25 helices, opposite where complexin binds^14,16^. The primary interface juxtaposes the membrane, where Syt-1 triggers fusion in the presence of calcium ions by disrupting the membrane and promoting the trans-to-cis transition of the SNARE complex^16^. As fusion proceeds, the vesicle membrane collapses into the plasma membrane, and the transmembrane domains of Syb-2 and Stx-1 interact, marking the transition to the cis-SNARE complex^9^. The SNAP-47—Stx-3—Syb-2 complex is thought to function via similar mechanisms, as both complexin and Syt-1 have been shown to regulate this complex, although synaptotagmin-7 (Syt-7) is functionally redundant with Syt-1 in this system^17^. Though immense progress has been made in understanding the regulation of membrane fusion during neurotransmitter release, dramatically less is known about the postsynaptic machinery used in AMPAR insertion, in large part due to a lack of structural information about SNAP-47.

SNAP-47 is particularly interesting due to its unique size and versatility as both a synaptic and an autophagic protein. Additionally, while human Stx-3 and Stx-1a have 62% sequence identity, SNAP-47 and SNAP-25 only have 12% identity. SNAP-47 is almost twice the molecular weight of SNAP-25, and in vitro fusion experiments have shown that SNAP-47-containing SNARE complexes have slower kinetics than SNAP-25-containing complexes^18^. However, it is unclear what underlies these kinetic differences. One possible explanation for the differing kinetics relates to differences in membrane tethering. SNAP-47 lacks the poly-cysteine region found in SNAP-25, which allows for SNAP-25 to be palmitoylated and inserted into the plasma membrane^12^. However, SNAP-47 does contain one predicted Pleckstrin homology (PH) domain, which can associate with phosphoinositides and is important for proper localization of SNAP-47 during autophagy^19,20^. Another possibility relates to SNAP-47’s post-translational modifications. Acetylation of three lysines within SNAP-47 has been shown to impair autophagy mediated by SNAP-47—Stx17—VAMP7/8 SNARE complexes in HeLa/Parkin cells^21^. However, it has yet to be determined if acetylation of these lysines would impact SNARE complexes containing Syb-2 and Stx-3. Finally, SNAP-47 contains a ∼175-amino-acid region of no known function between its two SNARE domains, allowing for possible novel regulatory mechanisms.

Computational structural biology approaches have been used to substantially advance the understanding of SNARE-mediated membrane fusion. For example, recent molecular dynamics (MD) studies have simulated SNARE-mediated plasma membrane and vesicle fusion events with atomistic detail^22^. Despite its utility, MD is computationally demanding and is not well-suited for identifying conformational changes that occur on the millisecond timescale or for simulating systems with large numbers of atoms. Recently, BioEmu, a deep-learning model that applies a generative diffusion model sampling an approximate equilibrium ensemble trained on MD trajectories and AlphaFold structures, has been developed to efficiently emulate the conformational landscape of proteins^23^. Because BioEmu is trained to emulate the equilibrium distribution directly rather than propagate dynamics through time, it bypasses the prohibitive simulation times an MD trajectory would need to sample millisecond-scale conformational changes. Additionally, the structure similarity tool Foldseek has allowed for the analysis of complex datasets, including those generated by BioEmu^24^. Together, these new tools exemplify the evolving suite of resource-efficient approaches that can supplement resource-intensive MD simulations.

To better understand the molecular mechanisms of SNAP-47-mediated membrane fusion and the effect of post-translational acetylation on SNARE complex structure, we developed a computational workflow combining MD and artificial intelligence tools, including AlphaFold3 (AF3)^25^, BioEmu, and Foldseek, and validated structural insights with biochemical assays. Using AF3, circular dichroism spectroscopy, and lipid-binding assays, we identified a second PH domain (PH2), and BioEmu and MD simulations showed that the two PH domains form a stable interface with each other and with the plasma membrane that helps orient synaptotagmin’s binding site towards the plasma membrane, thereby potentially facilitating fusion. PH1 forms a stable interface with the SNARE, and in silico acetylation of a key lysine in the PH1—SNARE interface disrupts this interface, untethering the PH domains from the SNARE complex. Together, these findings demonstrate how two previously poorly characterized PH domains orient the SNARE complex involved in LTP.

## Results

### AlphaFold3 identifies multiple SNAP-47—Stx-3—Syb-2 SNARE conformations

To characterize the architecture of the SNARE complex involved in LTP, we first used AlphaFold3 to predict 100 ternary complexes containing full-length SNAP-47 and the SNARE domains of Stx-3 and Syb-2 (Fig. 1A). Interface predicted template modeling (ipTM) values ranged from 0.78 to 0.90 and predicted template modeling (pTM) values ranged from 0.48 to 0.51 for the predicted complexes, with predicted local distance difference test (pLDDT) values consistently greater than 70 for predicted domains and less than 50 for large linker regions (Supplementary Fig. 1A). The canonical coiled-coil structure of the SNARE bundle was present in all conformations, but surprisingly, all predictions also contained two Pleckstrin homology/Pleckstrin homology-like domains (both referred to as PH domains) within SNAP-47. The first PH domain was found at the N-terminus (PH1), while the second (previously unidentified) PH domain (PH2) was found between SNAP-47’s two SNARE domains (Fig. 1B). To confirm our observation, we used Foldseek to search the CATH50 database for three-dimensional structural alignment with solved structures. For PH1, all ten top hits belonged to the Pleckstrin homology domain (PH domain)/Phosphotyrosine-binding domain (PTB) superfamily, and the query coverage ranged from 94.1% to 97.0%. For PH2, nine of the top ten hits were from the PH superfamily, with coverage ranging from 80.7% to 98.2%. Hierarchical clustering of aligned AlphaFold3 results via an all-versus-all, root-mean-square-deviation (RMSD) analysis revealed two main conformations: a “bound” (54% of predicted structures) and an “unbound” (40%) conformation defined by whether the PH domains were bound to one another (Supplementary Fig. 1B). The bound conformation was characterized by a ∼1200 Å^2^ buried surface area between PH1 and PH2 (Fig. 1B, top row and Supplementary Fig. 1C-D), while this interface was absent in the unbound conformations.

**Figure 1.**
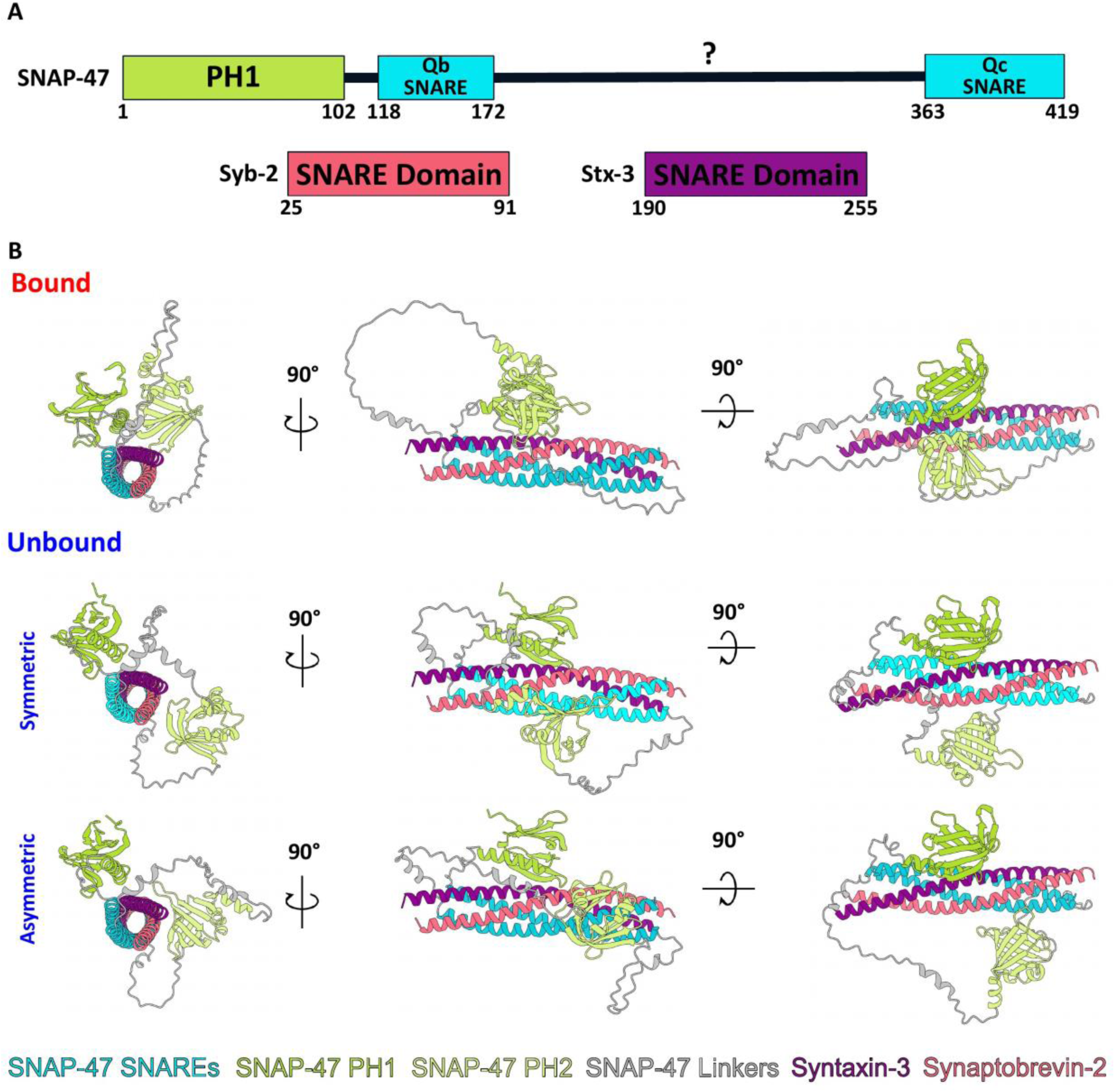
AlphaFold3 identifies multiple SNAP-47—Stx-3—Syb-2 SNARE conformations. **(A)** Domain maps for SNAP-47, Syb-2, and Stx-3. **(B)** AlphaFold predicted one bound conformation and two unbound conformations, asymmetric and symmetric. Structures are shown in three different views at 90-degree rotations from each other.

Within the unbound classes, we identified a continuum between symmetric and asymmetric subtypes, where each subtype was defined by the positioning of PH2. In the symmetric unbound conformations, PH1 and PH2 were at similar locations along the axis running through the SNARE bundle, but PH2 was located ∼140° rotated around the SNARE coiled coil from PH1 (Fig. 1B, middle row). Meanwhile, in the asymmetric unbound conformation, PH2 was translated down the coiled coil and at ∼120° rotation from PH1 (Fig. 1B, bottom row). Together, in the unbound conformation, PH2 sampled Syb-2 and SNAP-47’s C-terminal SNARE domain (Qc-SNARE); however, in the bound conformation, PH2 formed an interface with the SNARE bundle via interactions with Stx-3 and Syb-2 (Supplementary Fig. 1C). In all conformations, PH1 formed a ∼600 Å^2^ buried surface area with Stx-3, though the exact interface was subtly shifted between the bound and unbound conformations (Supplementary Fig. 1E).

### SNAP-47 PH domains contain antiparallel β-sheets and bind lipids

To validate the structural predictions made by AlphaFold3, we purified recombinant full-length mouse (*Mus musculus*) SNAP-47 (Supplementary Fig. 2A) and performed CD spectroscopy. The spectrum of purified SNAP-47 contained signal characteristic of a protein containing both α-helices and β-sheets (Fig. 2A). BeStSel^26^ spectrum analysis of the experimental spectrum revealed that SNAP-47 contained 31.4% α-helices, 17.7% β-sheets (14.0% antiparallel and 3.7% parallel), and 16.5% turns, consistent with the presence of multiple antiparallel, β-sheet-containing domains, like PH domains, and consistent with the predicted spectrum for mouse SNAP-47. We then purified and characterized human (*Homo sapiens)* PH1 alone and found 26.1% α-helices, 12.5% antiparallel β-sheets, and 15.9% turns (Fig. 2B, Supplementary Fig. 2B). We also purified and characterized human PH2 alone and found 0.8% α-helices, 39.7% β-sheets (36.1% antiparallel and 3.6% parallel), and 13.7% turns (Fig. 2C, Supplementary Fig. 2C). Therefore, the CD spectra showed antiparallel β-sheets throughout SNAP-47 and specifically within the PH1 and PH2 regions.

**Figure 2.**
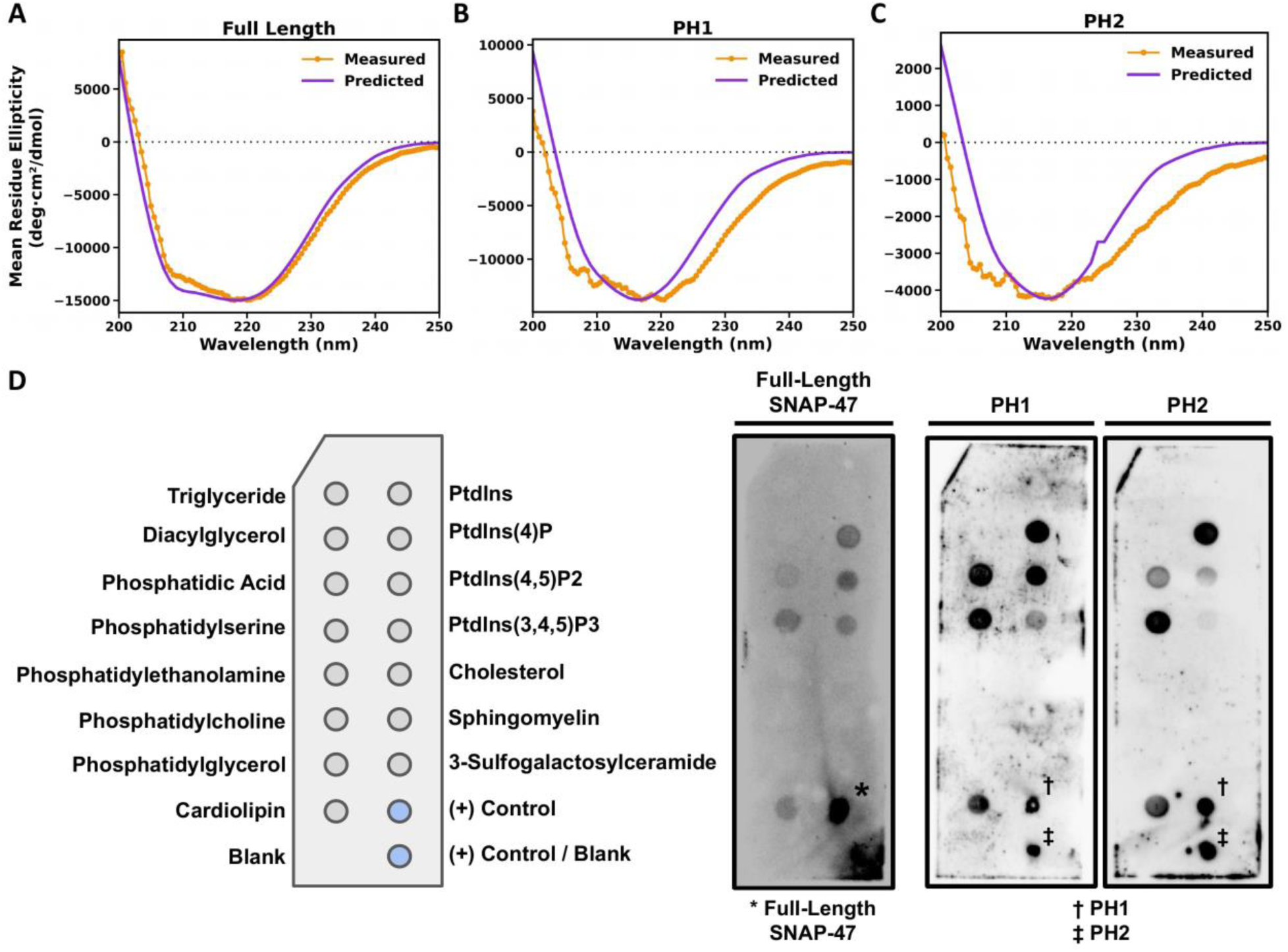
Structural characterization and lipid-binding properties of SNAP-47 PH domains. **(A)** CD spectrum of purified mouse SNAP-47 compared to the predicted mouse SNAP-47 spectrum. **(B)** CD spectrum of purified human PH1 compared to the predicted human PH1 spectrum. **(C)** CD spectrum of purified human PH2 compared to the predicted human PH2 spectrum. **(D)** Lipid-binding assay of purified full-length mouse SNAP-47, and human PH1 and PH2 domains.

To assess whether the predicted PH domains could associate with membrane lipids, we performed a lipid strip binding assay. Full-length SNAP-47 associated with several phosphatidylinositol (PI)-containing lipids, including PI(4)P, PI(4,5)P2, and PI(3,4,5)P3. In addition, it also associated with several anionic phospholipids, including phosphatidylserine (PS), cardiolipin (CL), and phosphatidic acid (PA). To test whether both PH domains could bind lipids, we performed the assay on isolated PH1 and PH2 domains and found that both domains retained lipid-binding capabilities (Fig. 2D). These domains therefore possess both structural and functional characteristics expected of PH domains.

To better characterize PH1 and PH2 lipid binding, we performed molecular docking of the PIP2 headgroup to each domain and identified favorable binding cavities within both PH1 and PH2 (Supplementary Fig. 3A). Following identification of favorable sites, we performed four randomly seeded, 1-µs MD simulations for each domain in the presence of a PIP2-containing lipid bilayer in GROMACS^27,28^ using the CHARMM36m force field^29,30^. Consistent with previous experimental results, MM/PBSA analysis of the interface between PH1 and the lipid bilayer revealed that three residues within the canonical PH lipid-binding β1-β2 loop contributed to PIP2 binding: K19, R20, and R21 (Supplementary Fig. 3B-C)^19^. Additionally, R84 demonstrated PIP2 specificity. PH2 lacked the canonical binding motif but was still able to associate with the membrane via binding to PS-DHA, a phosphatidylserine species enriched in the brain (Supplementary Fig. 3B, D).

### Molecular dynamics simulations and BioEmu reveal preference towards PH1—PH2 bound conformation

To test the stability of the AF3 predictions and gather insight into residue-specific interactions, we performed four randomly seeded, 1-µs MD simulations on each of the three conformations identified by AF3. Global descriptive metrics revealed the plateauing of RMSD at ∼1 nm for all conformations and the relative stability of residues within the SNARE and PH domains, while linker regions within SNAP-47 exhibited dramatic fluctuation (Supplementary Fig. 4, 5). In the bound conformation, the PH domains occupied a consistent surface across the length of the simulation in all four seeds (Fig. 3A, top row, and Supplementary Fig. 6). However, in both the unbound conformations, we observed dramatic movements of the PH2 domain (Fig. 3A, bottom two rows, and Supplementary Fig. 6). Overlaying the endpoint consensus map of each conformation with the other conformations revealed substantial overlap in the unbound structures (Fig. 3B), suggesting interconversion of the two unbound conformations on ns to µs timescales. To quantify the movement of PH2, we defined a polar axis through the SNARE complex and plotted domain positioning relative to PH1. Indeed, PH2 moved substantially in the unbound conformations and sampled the same region, while there was minimal movement in the bound conformation at its unique PH2 binding site (Fig. 3C).

**Figure 3.**
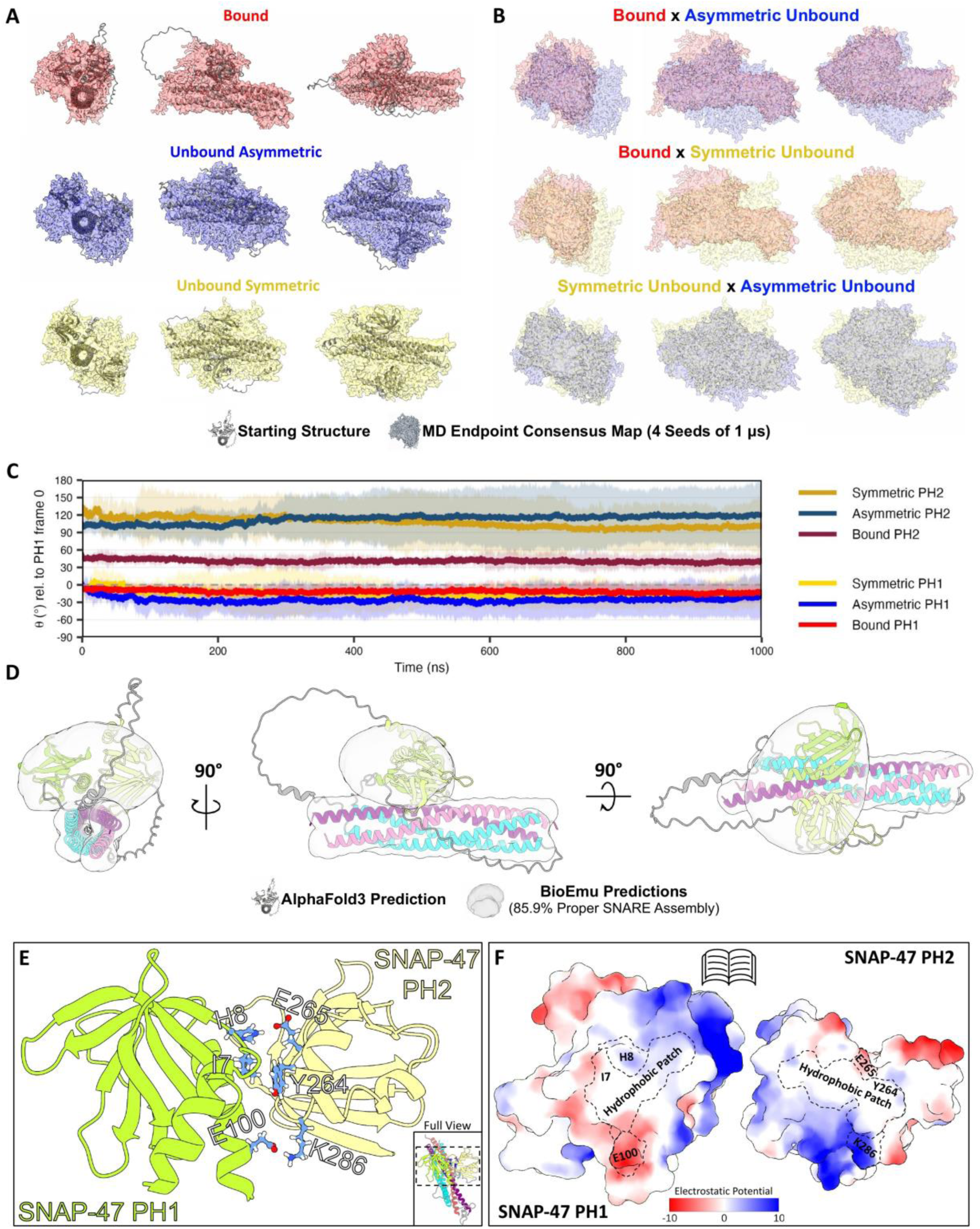
Molecular dynamics and BioEmu identify dynamic PH2-domain movement and a predominant bound state. **(A)** Overlay of each conformation’s starting structure versus the last frame of each trajectory (4 random seeds per conformation). Bound in red, unbound asymmetric in blue, and unbound symmetric in yellow. Structures are shown in three different views at 90-degree rotations from each other. **(B)** Overlay of each conformation’s combined final frames compared to each other. **(C)** Graph showing PH domain angle normalized to PH1 at t=0 for each conformation. Line is the average, while fainter background colors are 95% confidence intervals. **(D)** BioEmu predictions shown as a gray voxel map of average occupancy. Voxel map is overlaid on the bound AlphaFold3 prediction. Structures are shown in three different views at 90-degree rotations from each other. **(E)** Highest energy contributing residue pairs present in most simulations contributing to PH1—PH2 interface stability, shown with atomistic detail. **(F)** Open-book view of PH1—PH2 interface as an electrostatic map. Residues from panel E are labeled as patches.

MD demonstrated that the unbound conformations could interconvert, so next we asked whether the bound and unbound conformations could interconvert. Initially, we extended one of the unbound seeds to 2 µs but did not observe conversion to the bound conformations (Supplementary Fig. 7). We hypothesized that conversion between the bound and unbound conformations happens on a timescale that cannot be readily observed by MD. Therefore, we used BioEmu to generate 10,000 conformations of the ternary complex, emulating the equilibrium distribution. Because BioEmu is currently only compatible with single chains, we linked SNAP-47 to Syb-2 and Stx-3 with a GSG repeat. To ensure linker length was not disrupting complex formation, we tested 6-, 12-, 18-, 24-, and 30-amino-acid linkers and found that the 18-amino-acid linkers led to the most stable complex formation. We used Foldseek to filter out structures in which the coiled-coil assembly failed to form and found that, in the 18-amino-acid linker structures, 85.9% of predictions contained stable SNARE complexes (compared to 4.0%, 28.3%, 66.2%, and 50.9% for the 6-, 12-, 24-, and 30-amino-acid linkers, respectively) (Supplementary Fig. 8). We therefore chose to use an 18-amino-acid linker to minimize the likelihood of linker-induced artifact. A surface model generated from these structures confirmed that the SNARE complex was well resolved and that the PH domains frequently occupied the same region as seen in the bound conformation (Fig. 3D). Furthermore, 74.5% of the predictions with a well-defined SNARE complex contained a PH1—PH2 interface (Supplementary Fig. 9A), with PH1 and PH2 typically being offset by ∼55° (Supplementary Fig. 9B), consistent with the bound conformation identified by AlphaFold3.

As the bound conformation emerged as a more prominent conformation, we next decided to better characterize the PH1—PH2 interface that defined it. We observed that the PH1—PH2 interface persisted in all four bound seeds through the entirety of the 1-µs simulation (Supplementary Fig. 10A) and was stable even when one simulation was extended to 3 µs (Supplementary Fig. 11). To identify residues that drive the stability of the bound conformation, we performed MM/PBSA analysis on the MD trajectories, which revealed that the PH1—PH2 interface is maintained by H8-E265 hydrogen bonding, an E100-K286 salt bridge and several hydrophobic interactions, including a stronger I7-Y264 interaction, within the core of the interface (Fig. 3E-F, Supplementary Fig. 10B)^31^. Umbrella sampling was used to approximate binding free energy, which we present as ΔG_PMF_, the difference between the minimal energy state of the complex and the dissociated state. This revealed that the ΔG_PMF_ for the highest-scoring MM/PBSA seed was -13.9 kcal/mol (Supplementary Fig. 12A). Therefore, MD and BioEmu identified the PH1—PH2 interface as biologically plausible.

### PH1—SNARE form a stable interface that is disturbed by acetylation

The PH1—SNARE interface was consistent across all AF3 predictions, so we decided to further investigate this interface. Umbrella sampling indicated that PH1 and the SNARE domains interacted with a ΔG_PMF_ of -8.5 kcal/mol (Supplementary Fig. 12B). Buried surface area analysis of the PH1—SNARE interface within the bound conformation showed that the PH1—SNARE interface is maintained mainly by PH1 interacting with the Qb-SNARE (N-terminal SNARE domain) of SNAP-47 (Supplementary Fig. 10C) and Stx-3 (Supplementary Fig. 10D), with transient interactions with Syb-2 (Supplementary Fig. 10E). MM/PBSA analyses identified several salt bridges within the core of the PH1—SNARE interface (Supplementary Fig. 10F). SNAP-47 forms an intramolecular E95-R132 salt bridge, and three PH1—Stx-3 salt bridges, including R99-D217, K58-E223, and K58-E227 (Fig. 4A-B).

**Figure 4.**
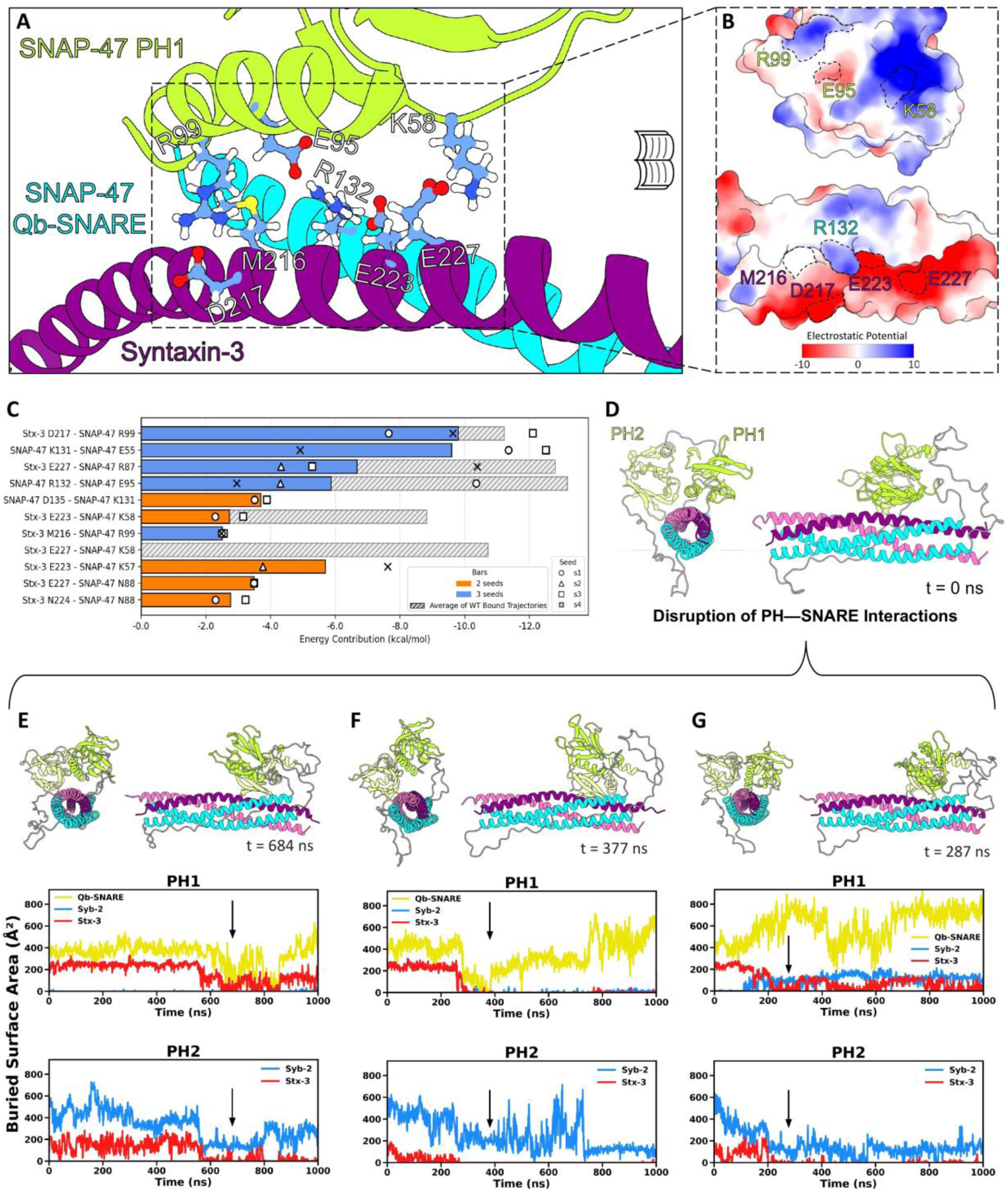
PH1—SNARE interface stability can be disrupted by acetylation. **(A)** Highest energy contributing residue pairs present in most simulations contributing to PH1—SNARE interface stability, shown with atomistic detail. **(B)** Open-book view of PH1—SNARE interface as an electrostatic map. Residues from panel A are labeled as patches. **(C)** MM/PBSA residue pair energy contributions in acetylation replicates (orange and blue bars, seed markers) compared to wild type (grey bars, averaged). **(D)** Starting SNAP-47 structure for acetylation runs. **(E-G)** Three acetylation MD replicates where the PH1—SNARE interface is disrupted. Quantification of the buried surface area for each seed is shown below. Arrows identify the time of the disruption event.

SNAP-47 possesses endogenous acetylation sites that disrupt autophagosome function^21^. This acetylation occurs at K58, which we identified as a notable energetic contributor to the PH1—SNARE interface via its interactions with E223 and E227 on Stx-3. To probe whether an acetylation event at the interface between PH1 and the SNARE complex impacts the SNAP-47—Syb-2—Stx-3 SNARE complex formation, we first performed a K58Q mutation and generated 10,000 conformations using BioEmu, as BioEmu does not currently support post-translational modifications. In this acetylation mimic, we observed a 2.4-fold decrease in SNARE coiled-coil assembly (85.9% to 35.2% proper assembly) (Supplementary Fig. 13A), as well as an increase in PH domain spread across the coiled-coil barrel (Supplementary Fig. 13B), suggesting that acetylation destabilizes the complex. To confirm these BioEmu results, we acetylated K58 in silico and performed MD. Across four seeds, MM/PBSA unsurprisingly revealed a complete loss of the Stx-3 E227 and SNAP-47 K58 salt bridge (Fig. 4C). Additionally, two replicates saw the PH domains dissociate from the SNARE coiled-coil structure, an event we did not observe in any of the wild-type simulations (Fig. 4D-F). To test whether this disruption of the PH1—SNARE interface was reproducible, we ran four additional seeds. We observed one additional disruption event (Fig. 4G) and one instance of the PH domains rotating around the SNARE complex in a manner we had not observed before (Supplementary Fig. 14). Umbrella sampling was once more used to more accurately approximate the binding energy, where we observed that the acetylated structure had a ΔG_PMF_ of -6.6 kcal/mol between PH1 and the SNARE complex (Supplementary Fig. 12C), thus demonstrating that acetylation disrupts the interface between the SNARE and PH domains.

### PH2 dissociation permits complexin binding at its canonical site

We next sought to characterize the PH2—SNARE interface. Umbrella sampling found a ΔG_PMF_ of -8.1 kcal/mol between PH2 and the SNARE complex (Supplementary Fig. 12D). Buried area analysis of the PH2—SNARE interface revealed that PH2 transiently interacted with Stx-3 and formed a small, relatively stable interface with Syb-2, though in one seed (seed 4), PH2 temporarily completely dissociated from the SNARE complex (Supplementary Fig. 10G, H). Although PH2 only transiently interacted with Stx-3, MM/PBSA analysis revealed salt bridge formation between Stx-3 and PH2 consisting of D217-R229 (Fig. 5A).

**Figure 5.**
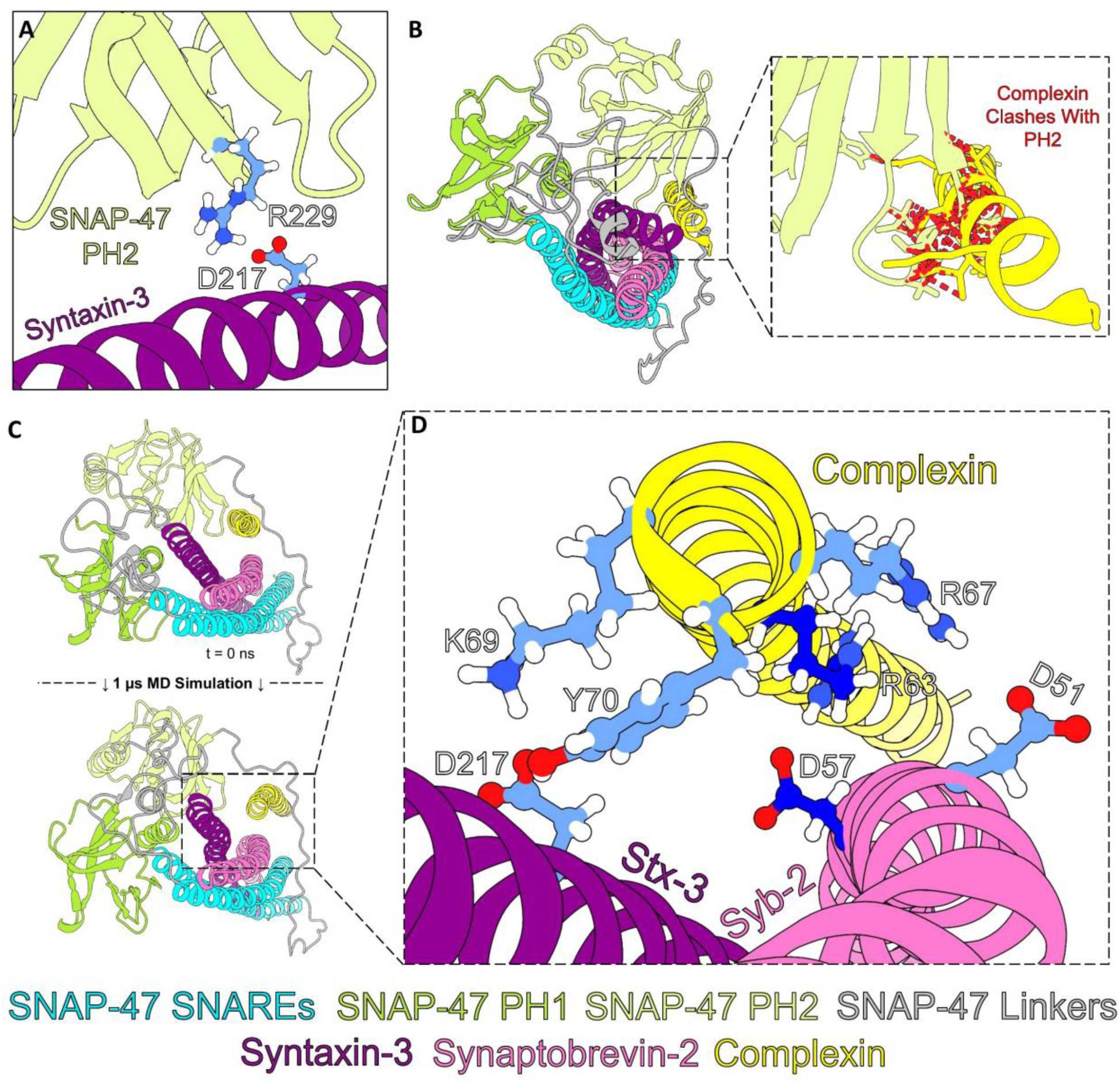
Complexin binds to canonical SNARE-binding site. **(A)** Highest energy contributing residue pair present in most simulations contributing to PH2—SNARE interface stability, shown with atomistic detail. **(B)** Overlay of a solved complexin (AA32-72) and SNARE coiled-coil structure over a relaxed bound conformation frame reveals that PH2 sterically clashes with complexin in its binding site. **(C)** Top structure shows the relaxed structure of complexin bound to the ternary structure with PH2 dissociated from its binding site. Bottom structure shows one of four seeds of MD, showing that complexin remains in the binding site and further dislocates PH2. **(D)** Magnification of panel C showing highest energy contributing residue pairs present in most simulations contributing to Complexin—SNARE interface stability, shown with atomistic detail.

Because complexin is a key regulator of this SNARE complex^32^, we tested whether the bound conformation is compatible with complexin binding. Initially, we observed that the acidic groove formed by Stx-1a and Syb-2, in which complexin binds, is maintained in Stx-3 and Syb-2 (Supplementary Fig. 15A). Therefore, we overlaid a solved crystal structure of complexin bound to the presynaptic SNAREs onto the SNAP-47-containing SNARE structure^13^ and observed that PH2, when associated with the SNAREs, occupies complexin’s binding site (Fig. 5B). To test whether PH2 would interact with complexin and whether complexin could be competed off the SNARE complex in the presence of PH2, we chose a frame from seed 4 of the bound conformation simulation where PH2 dissociated from the SNAREs and docked complexin to its canonical binding site. MM/PBSA analyses of four 1-µs MD simulations revealed that complexin interacted favorably with Syb-2 and Stx-3 (Fig. 5C). This interaction was driven mainly by salt bridges, including D51-R67, D217-K69, and D57-R63, as well as weaker electrostatic interactions such as D217-Y70 (Fig. 5D, Supplementary Fig. 15B). Umbrella sampling of complexin to the SNARE revealed a strongly favorable interaction (ΔG_PMF_ = -17.5 kcal/mol) (Supplementary Fig. 12E). We next tested possible interactions between PH2 and complexin and measured a ΔG_PMF_ of only -3.4 kcal/mol with umbrella sampling (Supplementary Fig. 12F). Together, these findings demonstrate that complexin can bind to its canonical SNARE complex binding site in the absence of the PH2—SNARE interaction and that there is no observable PH2—complexin interaction.

### PH1, PH2, and Syt-1 can simultaneously interact with the plasma membrane

Having identified PH—lipid, PH—SNARE, and complexin—SNARE interactions, we next aimed to observe these interfaces in a system containing all these interfaces. We built a system containing the SNAP-47—Syb-2—Stx-3—complexin complex and a plasma membrane containing PIP2^22^. We oriented the PH domains towards the plasma membrane and observed that the Syt-1 primary interface, which is essential for calcium-triggered fusion, was also in close proximity to the plasma membrane. To investigate whether the Syt-1 primary interface was maintained in SNAP-47, we compared the electrostatic surfaces of SNAP-25 and SNAP-47, where we observed that the Qb-SNARE of SNAP-47 lacked the negative patch found in SNAP-25 (Supplementary Fig. 16A). To test whether Syt-1 could still bind to its primary interface on SNAP-47, we aligned the C2B domain of Syt-1 (the domain essential for triggering fusion) containing two calcium ions onto the quaternary complex, added Stx-3’s transmembrane domain (AA 256-289), embedded this transmembrane domain into a plasma membrane, and performed a 4-µs MD simulation. We observed that Syt-1, PH1, and PH2 simultaneously interacted with the plasma membrane (Fig. 6A, Supplementary Fig. 16B). PH2 associated with PIP2 in the membrane via a binding pocket consisting of R220, T221, R269, and R281 (Fig. 6B, Supplementary Fig. 16C-D). Interestingly, we found one of SNAP-47’s characterized acetylation sites (K259) within this interface (Supplementary Fig. 16E). PH1 also bound to PIP2, both via an S82 and R84 interaction as well as its canonical β1-β2 loop (Fig. 6C-D, Supplementary Fig. 16F). Additionally, Syt-1’s C2B domain interacted with PIP2 with a binding pocket consisting of K321, R322, and K354 (Fig. 6E, Supplementary Fig. 16G), which are all residues within the polybasic region known to bind lipids^33^. Within this complex, Syt-1 also interacted with SNAP-47’s SNARE and maintained a consistent interface throughout the course of the entire simulation (Supplementary Fig. 16H-I).

**Figure 6.**
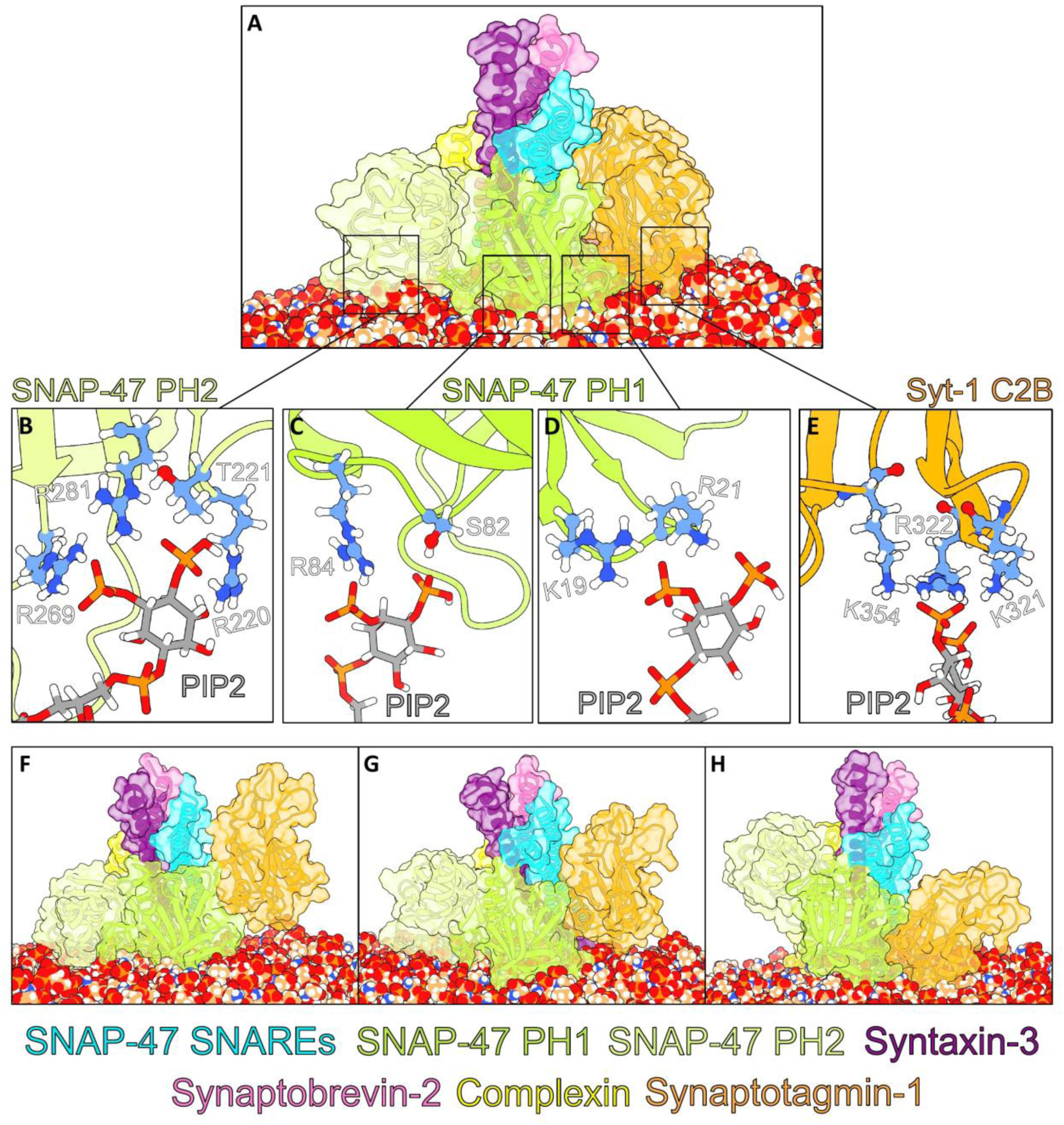
PH1 can orient synaptotagmin-1 towards the plasma membrane. **(A)** MD simulations showing PH1, PH2, and Syt-1 simultaneously binding to the plasma membrane. **(B)** Enlarged view of PH2—Membrane interface from panel A showing PH2’s PIP2 binding site with top-scoring residues shown in atomistic detail. **(C)** Enlarged view of PH1—Membrane interface from panel A showing PH1 bind PIP2 via a non-canonical binding site, with residues shown in atomistic detail. **(D)** Enlarged view of PH1—Membrane interface from panel A showing PH1 bind PIP2 via its canonical β1-β2 loop, with residues shown in atomistic detail. **(E)** Enlarged view of Syt-1—Membrane interface from panel A showing C2B’s polybasic region binding to PIP2. **(F)** MD simulation showing PH1 and PH2 binding to the plasma membrane, while Syt-1 remains unbound. **(G)** MD simulation showing PH1, PH2, and Syt-1 simultaneously binding to the plasma membrane, similar to panel A. **(H)** MD simulation showing PH1 and Syt-1 simultaneously binding to the plasma membrane while PH2 remains unbound.

To more thoroughly sample potential conformations, we performed three additional 1-µs simulations. In one seed, PH1 and PH2 interacted with the plasma membrane while Syt-1 did not associate (Fig. 6F, Supplementary Fig. 17A). Another seed replicated the long simulation where PH1, PH2, and Syt-1 simultaneously interacted with the membrane (Fig. 6G, Supplementary Fig. 17B). In the final seed, PH1 and Syt-1 interacted with the membrane while PH2 did not associate (Fig. 6H, Supplementary Fig. 17C). Therefore, we observed that PH1, PH2, and Syt-1 can simultaneously bind to the plasma membrane, though they can also adopt a conformation where one domain is separated from the membrane.

## Discussion

Here, we uncovered key structural differences between SNAP-25- and SNAP-47-containing SNARE complexes. SNAP-25 contains only its two SNARE domains, while SNAP-47 contains two PH domains in addition to its two SNARE domains. Using AF3, we identified a novel PH domain in SNAP-47 and validated these results with CD spectra collected from purified full-length mouse SNAP-47, human PH1, and human PH2, which confirmed the presence of β-sheets, consistent with the predicted PH domains. We functionally validated these domains by showing that they bind lipids. This finding is consistent with recent work that showed that even when SNAP-47 is truncated to lack PH1, it retains some lipid-binding capacity^20^. To our knowledge, this is the only SNARE protein with validated PH domains.

We next used MD, BioEmu, and umbrella sampling to characterize the architecture of the SNAP-47—Stx-3—Syb-2 complex. Three conformations emerged from the AF3 predictions: one bound state, where the two PH domains formed an interface, and two unbound states lacking the PH1—PH2 interface. Molecular dynamics simulations collapsed the two unbound conformations into one, and BioEmu predictions identified the bound conformation as the preferred orientation of the ternary complex. MD and umbrella sampling also showed the stability and energetic favorability of the PH1—PH2 interface within the bound conformation, which suggests that once the two PH domains meet, they remain bound together. This PH—PH domain interaction is consistent with pleckstrin behavior (for which the PH superfamily is named). Pleckstrin forms a dimer in its crystal asymmetric unit^34^ and in crosslinking experiments^35^.

In addition to the PH1—PH2 interface, we also observed a PH1—SNARE interface that is largely maintained by salt bridges. This mirrors the PH—coiled-coil interaction seen in the integrin-signaling adaptor, Skap-hom, which is also rich in salt bridges^36^. Interestingly, we found K58, an acetylation site in SNAP-47 known for regulating fusion, within the PH1—SNARE interface. Experimental data have shown that the acetylation of K58 disrupts autophagy, but whether this would structurally impact the LTP SNARE complex was untested. In this case, we first used BioEmu, instead of computationally demanding MD simulations, and found that the K58Q mutation destabilized SNARE complex formation. We then followed up with MD and performed in silico acetylation of this residue. We saw that not only was the salt bridge between Stx-3 E227 and SNAP-47 K58 disrupted, but also, in several simulations, PH1 completely dissociated from the SNARE. Notably, the dissociation events we observed were reversible, indicating that K58 acetylation does not completely ablate PH1—SNARE interactions but rather weakens them. Recent studies have found that deleting PH1 impacts PI4P-enhanced SNAP-47—Syb-2 interactions^20^, suggesting that disruption of the PH1—SNARE interface, such as by acetylation, might impair SNAP-47—Syb-2 association in the LTP SNARE complex. Though we did not directly test the effect of PH domain binding on the disassembly of SNARE complexes in this study, it is worth noting that αSNAP, a key adapter protein involved in SNARE disassembly, binds to SNAP-25 and Stx-1a in the cis-SNARE conformation at a site similar to the one PH1 occupies in the SNAP-47-containing complex^37^. Future work should explore the potential for PH domains (and the acetylation of PH1) to regulate SNARE complex disassembly.

Furthermore, we described a PH2—SNARE interface in which PH2 occupies complexin’s binding site, suggesting competition between PH2 and complexin for the groove between Stx-3 and Syb-2. Complexin and PH2 both bind to D217 on Stx-3, which was identified as the most energetically contributing residue within the PH2—SNARE interface. Because PH2 is intrinsic to the SNARE complex, we suggest a model where PH2 binds to Stx-3 D217, thereby occupying complexin’s binding site in the ternary complex, and must be displaced for complexin to bind. This competition is likely functionally relevant because postsynaptic knockdown of complexin impairs AMPAR insertion during LTP, and rescue requires complexin’s SNARE-binding activity^32^. Once complexin is bound, it is unlikely that PH2 continues to compete for complexin’s binding site due to the immensely higher binding affinity that complexin has for the SNAREs as suggested by umbrella sampling. Additionally, PH2 and complexin did not favorably interact as measured by umbrella sampling, reducing the likelihood of a recruitment role for PH2 in complexin binding.

When we combined the quaternary system with a plasma membrane and added the C2B domain of Syt-1, we observed PH1 playing a key scaffolding role in positioning the SNARE complex near the membrane. PH1 binds to both the SNARE bundle and PIP2, and upon membrane association, PH1 constrains the orientation of the SNARE bundle, keeping it near the plasma membrane. Importantly, this PH1-imposed orientation, which was maintained throughout the 4-µs simulation, positions the Syt-1 primary interface towards the plasma membrane, allowing C2B to simultaneously engage the SNARE bundle and PIP2 through its polybasic region. In the presynaptic complex, this represents the primed state of the complex, and our simulations suggest that the postsynaptic complex can adopt an analogous primed state^14,16,38^. This is notable because SNAP-47 lacks many of the negative residues found within Syt-1’s primary binding site on SNAP-25, raising the possibility that the primary interface would not be maintained. However, C2B remained engaged with PIP2 and the SNARE complex through the 4-µs simulation, suggesting that the scaffolding of the SNARE complex by PH1 may help compensate for the missing negative patch. Functionally, Syt-1 and Syt-7 perform seemingly redundant functions in LTP, and whether Syt-7 C2B can occupy the same site as Syt-1 C2B in this complex remains to be tested^17^. Finally, the dissociation of PH1 from the SNARE complex that we observed upon K58 acetylation would impair PH1’s ability to arrange the SNARE complex, perhaps impeding fusion.

Interestingly, a lipid blot revealed that PH2 also binds lipids, indicating that SNAP-47 engages the membranes through more than one anchor point. In further support of this, we observed PH1 and PH2 simultaneously interacting with the plasma membrane in three of four simulations. This is consistent with a recent study reporting that deletion of PH1 reduced, but did not abolish, PI4P-dependent membrane association of SNAP-47.^20^ Notably, the lipid preferences of PH2 were different when PH2 was isolated and when it was part of the full complex containing complexin and C2B. Experimental results on isolated PH2 indicated that PH2 can bind both PIP2 and PS. In simulation, isolated PH2 preferred PS-DHA over PIP2, whereas in the full complex PH2 preferred PIP2. The difference in binding preference between these two experiments can be explained by the fact that, as an isolated domain, PH2 had a wider range of orientations that it could sample the membrane with than it would in a complex. When bound to PH1 and the SNARE bundle, PH2 preferred PIP2 interaction. Interestingly, K259, a lysine experimentally determined to be acetylated, was consistently within the interface between PH2 and the membrane in our 4-µs simulation^21^. Acetylation at this position would neutralize a positive charge and could impede membrane association.

We identified two PH domains within SNAP-47 that can bind lipids. One possibility is that PH1 and PH2 contribute additively to membrane attachment as partially redundant attachment points (see dual anchoring model in Fig. 7A). Alternatively, the PH domains could bridge the plasma membrane and the membrane of AMPAR-containing vesicles, helping anchor these to fusion sites (see membrane bridging model in Fig. 7B)^39^. These models are not mutually exclusive, and a PH domain could transition from the AMPAR-containing vesicle to the plasma membrane prior to or during membrane fusion.

**Figure 7.**
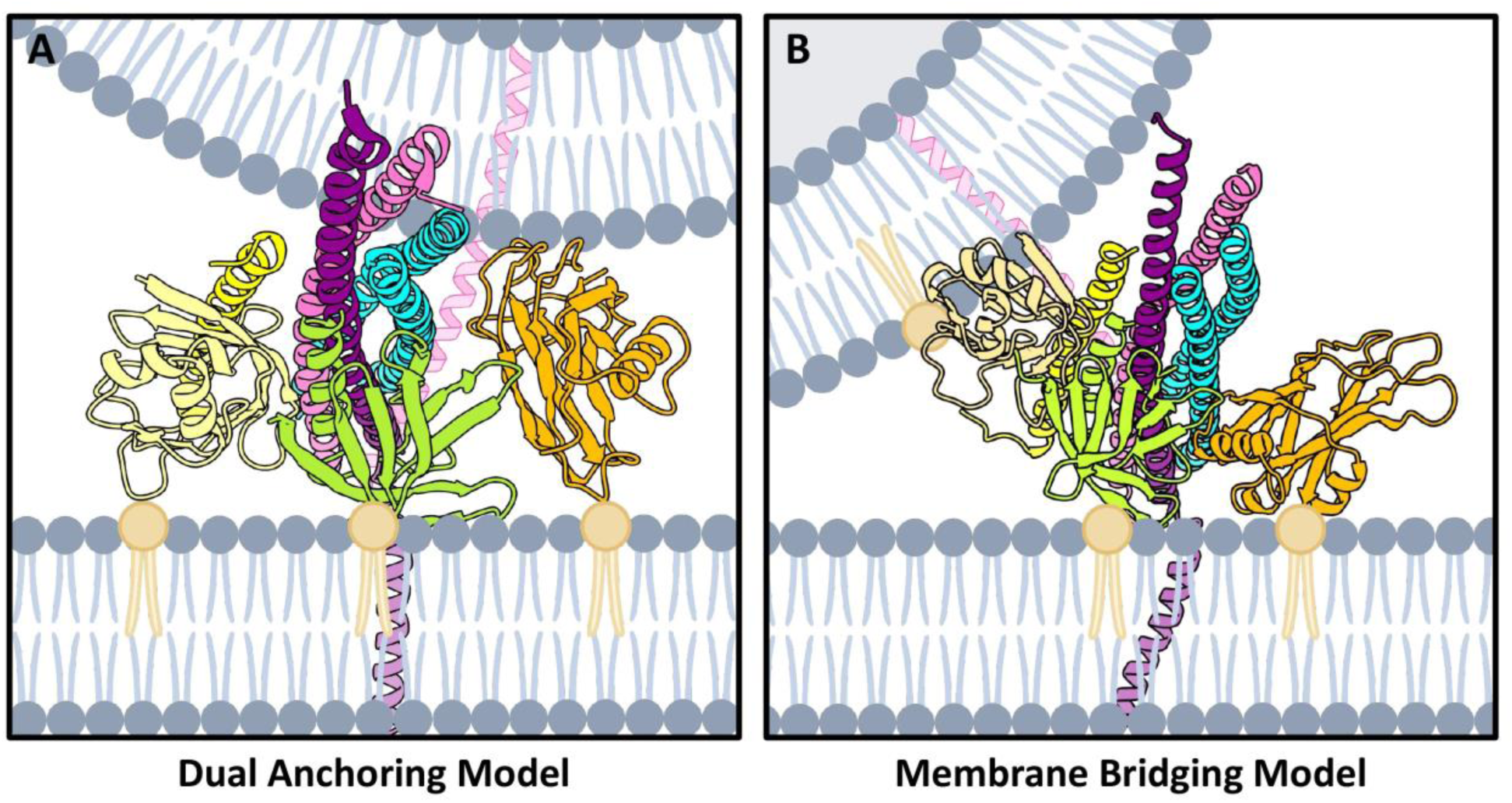
Summary model. **(A)** Dual anchoring model where PH1 and PH2 bind phosphatidylinositol-containing lipids (yellow lipid) in the plasma membrane, orienting Syt-1 towards the membrane, allowing it to also bind to the membrane. **(B)** In the membrane bridging model, one PH domain (PH1 shown here) and Syt-1 interact the same as in the dual anchoring model. The other PH domain (PH2 here) binds to phosphatidylinositol-containing lipids in the AMPAR-containing vesicle, serving as a tether between the membrane and the vesicle.

Here, we combined biochemical assays, MD, and BioEmu to characterize the structural and regulatory roles of SNAP-47’s PH domains in LTP SNARE-mediated membrane fusion. In sum, we propose that PH1 serves an organizing role, orienting the SNARE complex towards the plasma membrane in a way that facilitates Syt-1 binding to its primary interface. PH1 bound to the SNARE bundle and the plasma membrane (with PH2 dissociated from complexin’s binding site) is consistent with a pro-fusion state, while acetylated K58 inhibits PH1—SNARE engagement, inducing an anti-fusion state. This model provides a potential mechanistic framework for how post-translational modification and intramolecular domain interactions can regulate AMPAR delivery during LTP. We also presented a novel use case for MD and BioEmu in characterizing the architecture of protein complexes; future studies, including single-particle cryo-electron microscopy and single-molecule fusion assays, will be helpful for further testing our model. Together, these findings offer molecular insight into how neurons may use posttranslational modifications of SNARE proteins to modulate synaptic strength.

## Methods

### Sequences

Full-length human SNAP-47 (UniProt ID: Q5SQN1) and the SNARE domains of Syb-2 (AA 25-91) (UniProt ID: P63027) and Stx-3 (AA 190-255) (UniProt ID: Q13277) were used for AlphaFold screening, BioEmu emulations, initial ternary complex MD, and acetylation runs. The first 45 amino acids of SNAP-47 are from an alternative start codon. For complexin and Syt-1 experiments, we removed this unstructured region. All amino acid residue indices are reported with M46 having an index of 1 in alignment with UniProt ID: A0A087X0B7.

### AlphaFold3

AlphaFold3 predictions of the ternary complex were obtained by performing 100 randomly seeded prediction runs in the AlphaFold online server using the full-length SNAP-47 and the SNARE domains of Syb-2 and Stx-3. Predictions were aligned in ChimeraX using the matchmaker command, and RMSD between all pairwise combinations of structures was calculated. RMSD values were used as a distance metric for hierarchical clustering of structures via the unweighted pair group method with arithmetic mean (UPGMA).

### Recombinant protein purification

N-terminal 6x-His-tagged mouse SNAP-47 (UniProt ID: Q8R570), human PH1 (residues 6-104), or human PH2 (residues 202-314) (UniProt ID: A0A087X0B7) plasmid was transformed into BL21(DE3) cells and grown in 1 L cultures induced with 0.2 mM IPTG. Induced cells were incubated in an orbital shaker for 16 hours at 16 °C. Cells were lysed using a 1:1 ratio of B-PER (ThermoFisher) (4 mL per gram of cells), a buffer containing 50 mM Tris-Cl (pH 8.0), 600 mM NaCl, 1 mM TCEP, and 0.5% Triton X-100 (4 mL per gram of cells), Pierce protease inhibitor tablets (1 tablet per 50 mL buffer), lysozyme (1 mg/mL buffer), and DNase I (4 U/mL buffer). Lysate was solubilized in a buffer containing 8 M urea, 10 mM imidazole, 500 mM NaCl, 1 mM TCEP, and 50 mM Tris-Cl. Lysate was clarified at 18,800 x *g*, and protein was affinity-purified using Ni-NTA beads. Protein was refolded in 10% glycerol with a decreasing urea content from 8 M to a buffer containing no urea. Finally, protein was eluted with a buffer containing 10 mM sodium phosphate (pH 8.0), 1 mM TCEP, and 500 mM imidazole.

### Circular dichroism (CD) spectroscopy

Prior to CD spectra collection, protein was dialyzed overnight in 10 mM sodium phosphate (pH 8.0). CD spectra were collected on a Jasco J-720 Spectropolarimeter using Spectra Manager between 200 and 250 nm using a continuous scanning mode (100 nm/min scan speed, 0.5 nm data pitch, 1 s response time, 1.0 nm bandwidth, and 100 mdeg sensitivity). Data were averaged over 50 acquisitions to improve signal quality. High tension signal was kept under 800 V. BeStSel single spectrum analysis and fold recognition was used to assign the relative contribution of secondary structural elements to the collected spectrum. SESCA (v097) was used to generate a predicted spectrum for AF3-predicted structures.

### Lipid binding assay

Echelon Biosciences Membrane Lipid Strips were blocked for 1 h with TBST containing 3% fatty-acid-poor BSA before being incubated with purified PH1 or PH2 (5 μg/mL) for an additional 1 h. Following incubation, strips were washed with TBST and incubated with 6×-His Tag Monoclonal Antibody, HRP (1:900) for 1 h. Strips were then washed with TBST and developed using SuperSignal West Pico PLUS Chemiluminescent Substrate before imaging.

### BioEmu

BioEmu (v1.1.0) was used to generate a conformational ensemble comprising 10,000 independent structures of full-length SNAP-47 and the SNARE domains of Syb-2 and Stx-3. A glycine-serine-glycine (GSG) linker was added to facilitate *in silico* assembly of the SNARE complex. Five linker lengths were tested (6AA, 12AA, 18AA, 24AA, 30AA), and the 18-amino-acid (18AA) linker was selected based on its ability to produce stable predictions. To investigate the structural effects of post-translational modification, K58 was substituted with glutamine to mimic acetylation, and, again, 10,000 structural conformations were generated.

### Foldseek

The online Foldseek server was used to search the CATH50 database for three-dimensional structural alignment in 3Di/AA mode. Additionally, BioEmu predictions were clustered using Foldseek (release 10) with identical parameters across all analyses, including a TM-score threshold of 0.715 and a coverage cutoff of 0.90. Parameters were manually chosen to maximize the size of the largest class of structures from the wildtype prediction while filtering out structures with poorly assembled SNARE complexes. ChimeraX’s^40^ molmap command was used on aligned structures to generate density maps to visualize consensus structures.

### General MD

Starting structures for each conformation were selected by choosing a representative structure from the RMSD clustering results by randomly selecting a prediction from each cluster. Structures were solvated in a cubic box of SPC/E^41^ water with 0.15 M NaCl added to neutralize the net charge. Each system underwent energy minimization using the steepest descent algorithm until a maximum force of 1,000 kJ mol−1 nm−1 was reached, followed by 100 ps of NVT equilibration at 300 K with heavy atoms restrained. Next, 100 ps of NPT equilibration at 1 bar and 300 K was performed, also with heavy atom restraints. Production simulations were then carried out under unrestrained NPT conditions.

All simulations used the same force field and were performed in GROMACS (version 2022.2), except the Syt-1-containing simulations, which used version 2024.4. Temperature was maintained at 300 K using the velocity-rescale thermostat (τ = 1.0 ps), and pressure was controlled at 1 bar using the C-rescale barostat (τ = 5.0 ps).

Production simulations were run for a total of 1.0 μs per replicate unless otherwise noted. To more fully sample the potential energy surface, four randomly seeded replicates were generated for each conformation.

### Complexin MD

To determine clashing between PH2 and complexin, we used ChimeraX’s^40^ Clash feature with default settings. For MD simulations that included complexin, we manually selected frames from seed four of the bound conformation where MM/PBSA energies between PH2 and the SNAREs were weakest. We then used ChimeraX to align the AF3 full-length human complexin (UniProt Accession: O14810) prediction with the bound structure, using the solved Complexin/SNAP-25 structure as reference^13^. This structure was then used as the starting structure for the complexin runs. Runs were performed as in general MD. Production simulations were run for a total of 1.0 μs per replicate with four independent replicates. An extended equilibration protocol was employed to allow additional structural relaxation prior to production dynamics. Systems were solvated with SPC/E water and minimized using the same procedures described above, including steepest descent energy minimization under positional restraints. In contrast to the standard simulations, equilibration consisted of a 1-ns NVT phase followed by a 1-ns NPT phase, both performed with positional restraints maintained on the protein. Equilibration was carried out at 300 K using the velocity-rescale thermostat (τ = 1.0 ps), with pressure maintained at 1 bar during NPT equilibration using the C-rescale barostat (τ = 5.0 ps).

### PIP2 docking and lipid binding

The PH1 and PH2 domains were extracted separately from the bound conformation of the full-length protein, which was selected due to its structural stability and lowest RMSD observed in baseline simulations. Each domain was isolated as an independent structure using ChimeraX and prepared individually for docking and membrane simulations.

SwissDock’s Attracting Cavities 2.0^42–45^ was used for docking of PIP2 headgroup (SMILES: CCOP(O)(=O)O[C@@H]1[C@H](O)[C@H](O)[C@@H](OP(O)(O)=O)[C@H](OP(O)(O) =O)[C@H]1O) to the isolated PH1 and PH2 domains from the last frame of the bound conformation’s seed one trajectory. The docking search box was centered near PH1’s previously experimentally determined lipid binding site and included the entire membrane-exposed interface, while PH2’s box was centered to encompass the entirety of the domain for unbiased docking. Sampling exhaustivity was set to 5.

Protein–membrane systems were constructed using CHARMM-GUI’s Bilayer Builder^46,47^. Each PH domain was manually positioned approximately a few Å above the membrane bilayer as an initial separation prior to equilibration. The membrane was built in a rectangular box (100 Å × 100 Å) with a minimum water thickness of 10 Å on both sides and a lipid composition of 45% CHL1, 7% SAPE, 14% SDPE, 10% SDPS, 8% POPC, 5% SAPI2D, 10% SOPS, and 1% SAGL^38^. The system was solvated with 0.15 M KCl, and counterions were added to neutralize the total charge.

Equilibration followed the standard CHARMM-GUI protocol with multiple restrained steps, including two 125-ps NVT phases followed by four NPT phases (125–500 ps each) with gradually reduced positional restraints on protein and lipids. Production simulations were then performed under semi-isotropic NPT conditions at 300 K and 1 bar using the velocity-rescale thermostat (τ = 1.0 ps) and C-rescale barostat (τ = 5.0 ps). Production trajectories were generated using a 2-fs timestep for 500,000,000 steps (1.0 μs total simulation time), with coordinates written every 100 ps (50,000 steps). For each PH domain, four independent replicate simulations were performed. Membrane interaction energies were quantified using MM/PBSA calculations.

### Syt-1 MD

The final frame of seed 1 of the ternary complex in the presence of complexin was chosen as the starting structure, as it had the least steric clashing with synaptotagmin. The solved structure of Ca^2+^-bound Syt-1 (5CCG)^16^ was used to align Syt-1 with SNAP-47’s α-helices via MatchMaker, upon which Syt-1 was truncated to the C2B domain. Following this, the SNARE domain of Stx-3 was replaced with the AlphaFold3 prediction of the SNARE and transmembrane portion of Stx-3 via MatchMaker. Stx-3’s C-terminal region was then oriented to allow the transmembrane region of Stx-3 to be inserted into a membrane. This structure was used as the input for CHARMM-GUI.

CHARMM-GUI was then used to construct a bilayer with Stx-3’s transmembrane domain embedded and the PH1 PIP2 binding site oriented near the bilayer. The membrane was built in a rectangular box (150 Å × 150 Å) with a minimum water thickness of 20 Å. The same membrane composition and solvation were used as for the lipid binding membrane. All equilibration and minimization steps were performed as in the lipid binding membrane conditions except for the initial equilibration step, which had no atom constraints and a smaller emtol step to allow additional relaxation of clashes following membrane insertion.

### Acetylation MD

Following 1.0 μs of baseline MD simulations, the bound conformation was selected for acetylation studies, as it represented the most stable structure across replicates. The final frame of the lowest-RMSD replicate was used as the starting structure. Lysine acetylations were introduced as modified lysine residues using CHARMM-GUI’s PDB Reader & Manipulator^48–51^, with a single acetylation at K58. The modified structures were subsequently processed using CHARMM-GUI’s multicomponent assembler and exported in GROMACS format. Systems were solvated in a cubic box (∼170 Å per side) with TIP3P water and 0.15 M NaCl, with ions placed using a distance-based method.

Initial energy minimization was performed using the steepest descent algorithm for 5,000 steps with backbone and sidechain positional restraints (400/40 kJ/mol/nm²). Equilibration consisted of a 125-ps NVT phase (1 fs timestep) followed by a 125-ps NPT phase at 300 K and 1 bar, with positional restraints maintained. Production simulations were then performed under unrestrained NPT conditions using the same simulation parameters described above, including a 2-fs timestep, velocity-rescale thermostat (τ = 1.0 ps), C-rescale barostat (τ = 5.0 ps). Separate temperature coupling groups were used for solute and solvent. Four independent, 1.0-μs replicates were performed. Following observations in the initial four seeds, an additional four seeds were run for a total of eight seeds.

### Trajectory analysis

Completed MD trajectories were stripped of solvent using the built-in GROMACS tools. Trajectories were first visually inspected in ChimeraX. Root-mean-square deviation (RMSD) was plotted over the full trajectory, and mean RMSD values were calculated from the final 850 ns to avoid early equilibration behavior. Root-mean-square fluctuation (RMSF) was calculated in the same manner, except that it was done for each protein individually. The starting structure was used as the reference for both RMSD and RMSF and alignment was done via backbone atoms. To analyze interactions and binding affinities, we used the gmx_MMPBSA and gmx_MMPBSA_ana analysis packages^31,52^. MM/PBSA was performed per trajectory to obtain energy contributions. Values are reported as the mean ± standard deviation across four seeds.

To acquire more biologically realistic binding affinities between interfaces, we performed umbrella sampling to characterize the free-energy landscape associated with dissociation of interfaces. All interfaces of interest, except the acetylated PH1—SNARE interface, were trimmed from the final frame of the top-scoring 1-μs simulation, as determined by MM/PBSA, to retain only the domains of interest. For the acetylated structure, the starting pose used for all simulations was selected and trimmed down to the interfaces of interest. Umbrella sampling was performed along the center-of-mass distance between the domains. Steered MD was used to pull the interfaces away from each other, generating configurations spanning the bound-to-dissociated continuum. Configurations were extracted initially at 0.1-nm intervals and used as starting structures for independent umbrella windows. Windows were restrained using a harmonic potential with a force constant of 1000 kJ mol⁻¹ nm⁻² and simulated for 10 ns. Production trajectories were analyzed to obtain the distribution of the reaction coordinate for each window. All runs were performed at 310 K to emulate physiological conditions. The resulting distributions were combined using the GROMACS implementation of the weighted histogram analysis method (WHAM)^53^ to calculate the potential of mean force (PMF). Histogram overlap and PMF convergence were assessed to ensure adequate sampling. When gaps in WHAM were observed, additional windows were generated to ensure complete overlap. Binding free energy was determined as the difference between the minimum PMF value and the average value of the plateaued PMF following dissociation. The plateau was defined as the average PMF value of the last 20% of the pull distance. Statistical uncertainty was estimated by Bayesian bootstrap using the histogram resampling method (b-hist) with 200 bootstrap iterations. The resulting bootstrap mean profile and associated standard deviations were used to define the error bands. Binding energies obtained from umbrella sampling are referred to as ΔG_PMF_ even though standard-state correction was not applied. As the pull coordinate was defined as a one-dimensional center-of-mass distance along the z-axis with no restriction on lateral (xy) movement, the unbound protein was free to sample the full simulation box cross-section, making unambiguous definition of the area term in the standard-state correction impossible. Use of the full box cross-section produced corrections of the wrong sign, while the effective area approximation introduced dependence on the box length used for that system. As the solvated boxes used for each system were slightly different, the standard-state correction was omitted and raw PMF well depths are reported with the plateau region referenced to zero.

To quantify domain interactions across trajectories, we calculated the buried surface area between domain pairs using the Shrake-Rupley algorithm (1.4 Å probe radius) as implemented in MDTraj, where buried surface area equals the sum of the solvent accessible surface area (SASA) of each molecule alone minus the SASA of the complex^54^. To determine the radial position of the PH domains in relation to the SNARE bundle, we calculated each domain’s center of mass and projected it onto a radial coordinate frame defined by the coiled-coil barrel axis. A rotation angle, θ, was computed for PH1 and PH2 at each frame and was visualized using polar histograms.

### Statistics and reproducibility

All simulations were run as independent, randomly seeded replicates, totaling approximately 42 µs across 36 trajectories (not counting umbrella sampling). We did not apply inferential statistical tests because MD replicates are not independent draws from a population in the way assumed by significance tests. The number of replicates was determined by computational cost rather than by an *a priori* power calculation, and p-values calculated for two to eight trajectories would suggest a precision our approach does not support. Instead, we reported every replicate individually so that the variability between seeds is directly visible. Differences between conditions are described qualitatively, and we restrict mechanistic claims to observations seen across multiple seeds.

## Data availability

All data are available in the main text or supplementary figures. MD trajectories have been deposited in Zenodo (10.5281/zenodo.21366152).

## Acknowledgements

This work was supported by the National Science Foundation (Award Number: 2437522) and the University of Richmond’s Arts & Sciences Summer Fellowship Grant. Figure 7 was created using BioRender.com.

## Author Contributions

IA: conceptualization, data curation, formal analysis, validation, investigation, visualization, methodology, writing – original draft, and writing – review and editing. MW: data curation, formal analysis, investigation, visualization, and writing – review and editing. RT: formal analysis and investigation. CF: formal analysis, investigation, and visualization. LL: formal analysis and investigation. CB: formal analysis and writing – review and editing. ChC: investigation. CaC: investigation. LS: investigation and writing – original draft. JP: conceptualization, data curation, formal analysis, validation, investigation, visualization, methodology, writing – original draft, writing – review and editing, supervision, funding acquisition, and project administration.

---

**Supplementary Figure 1.**
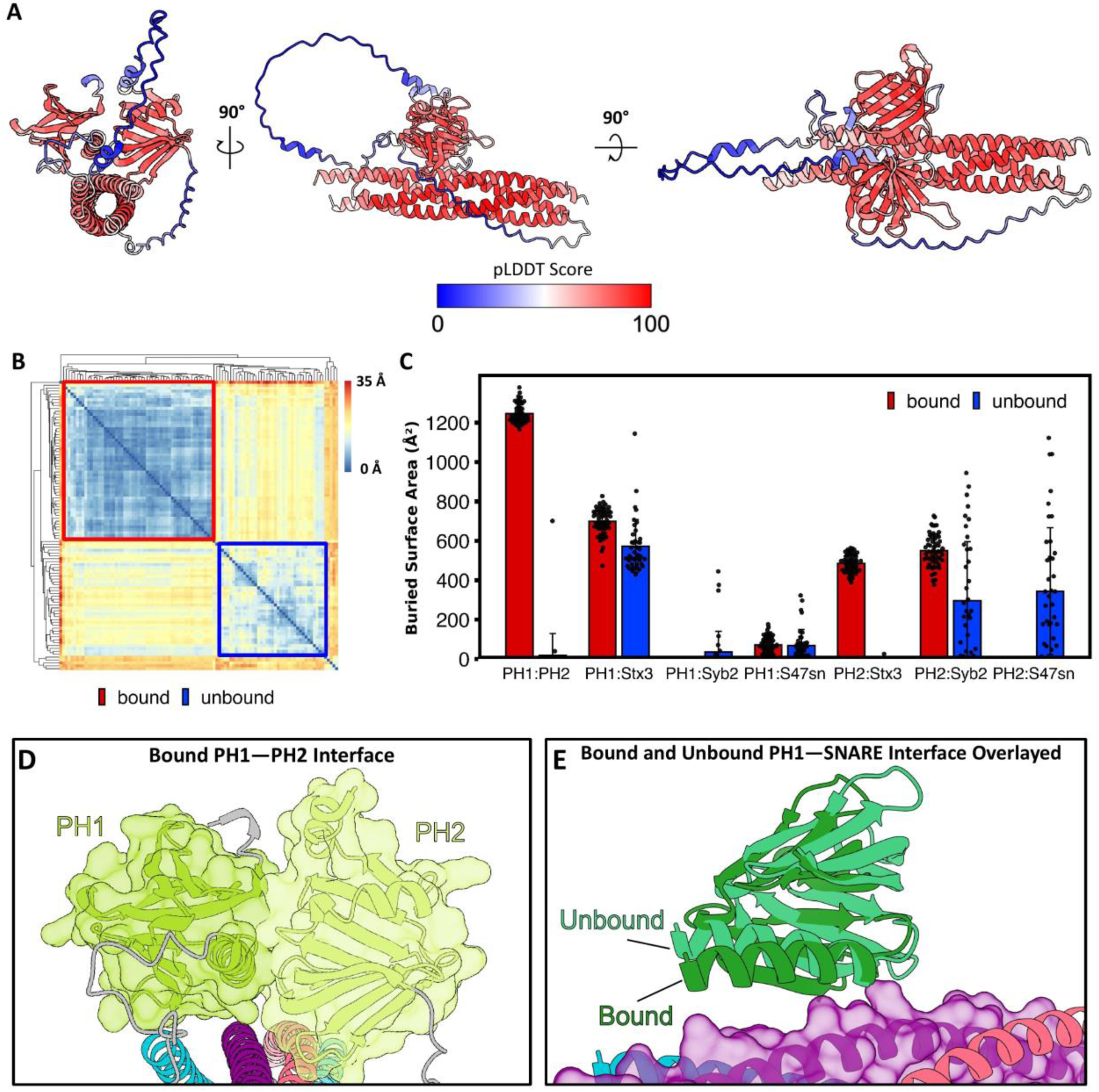
Characterization of AlphaFold3 ternary structure predictions. **(A)** Representative pLDDT of bound conformation. **(B)** Conformations were clustered via RMSD in an all vs. all comparison, with bound appearing in well-defined clusters outlined in red and unbound in a less defined cluster outlined in blue. **(C)** Interface area between SNAP-47’s PH domains and SNARE bundle protein domains, with bound in red and unbound in blue. **(D)** Space-filling model showing interface between SNAP-47’s PH domains. **(E)** Overlay of Bound and Asymmetric Unbound PH1—SNARE interfaces showing similarity in PH1 orientation.

**Supplementary Figure 2.**
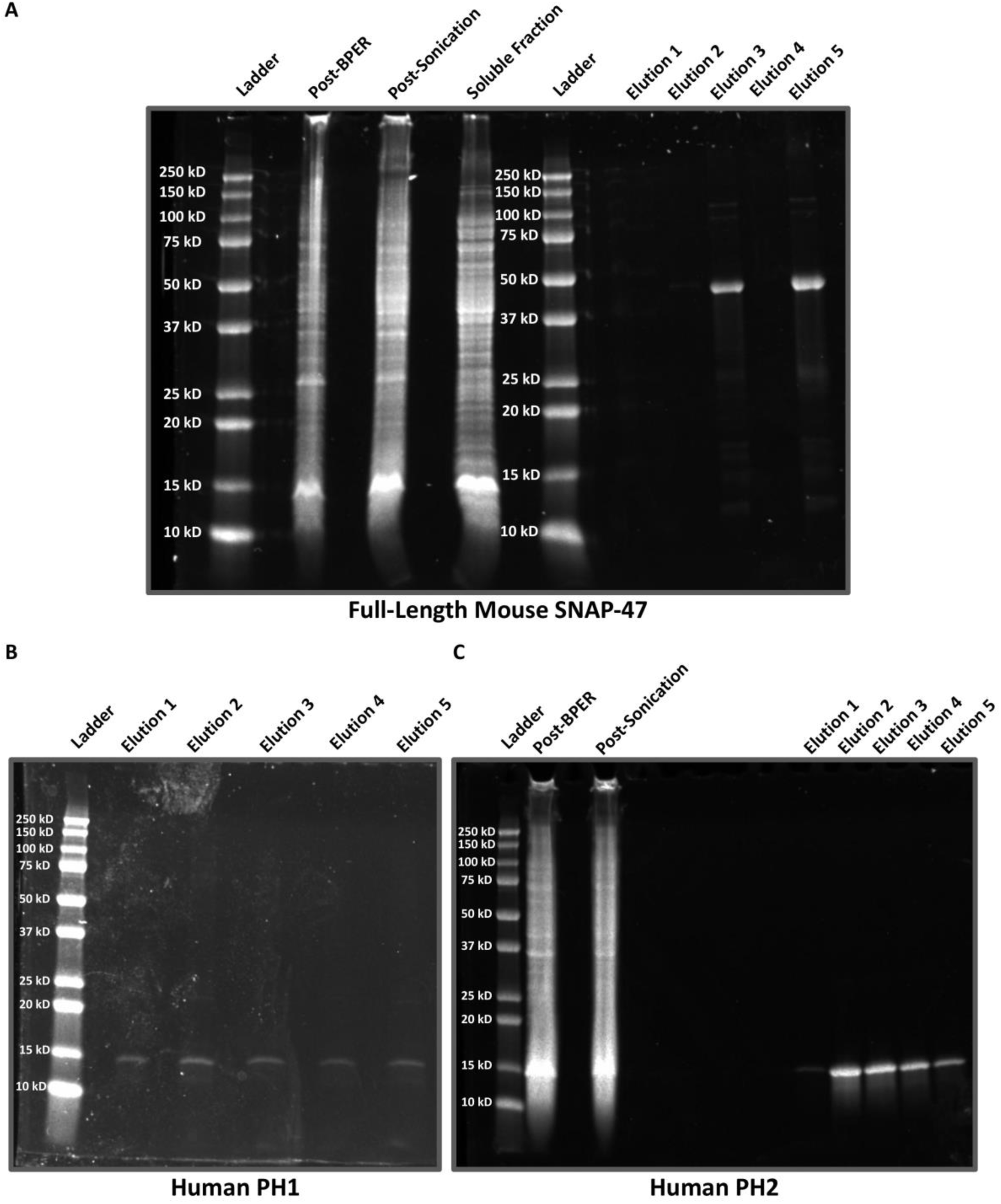
Purification of recombinant SNAP-47. **(A)** SDS-PAGE gel showing fractions from the purification of full-length mouse SNAP-47. **(B)** SDS-PAGE gel showing fractions from the purification of human PH1. **(C)** SDS-PAGE gel showing fractions from the purification of human PH2.

**Supplementary Figure 3.**
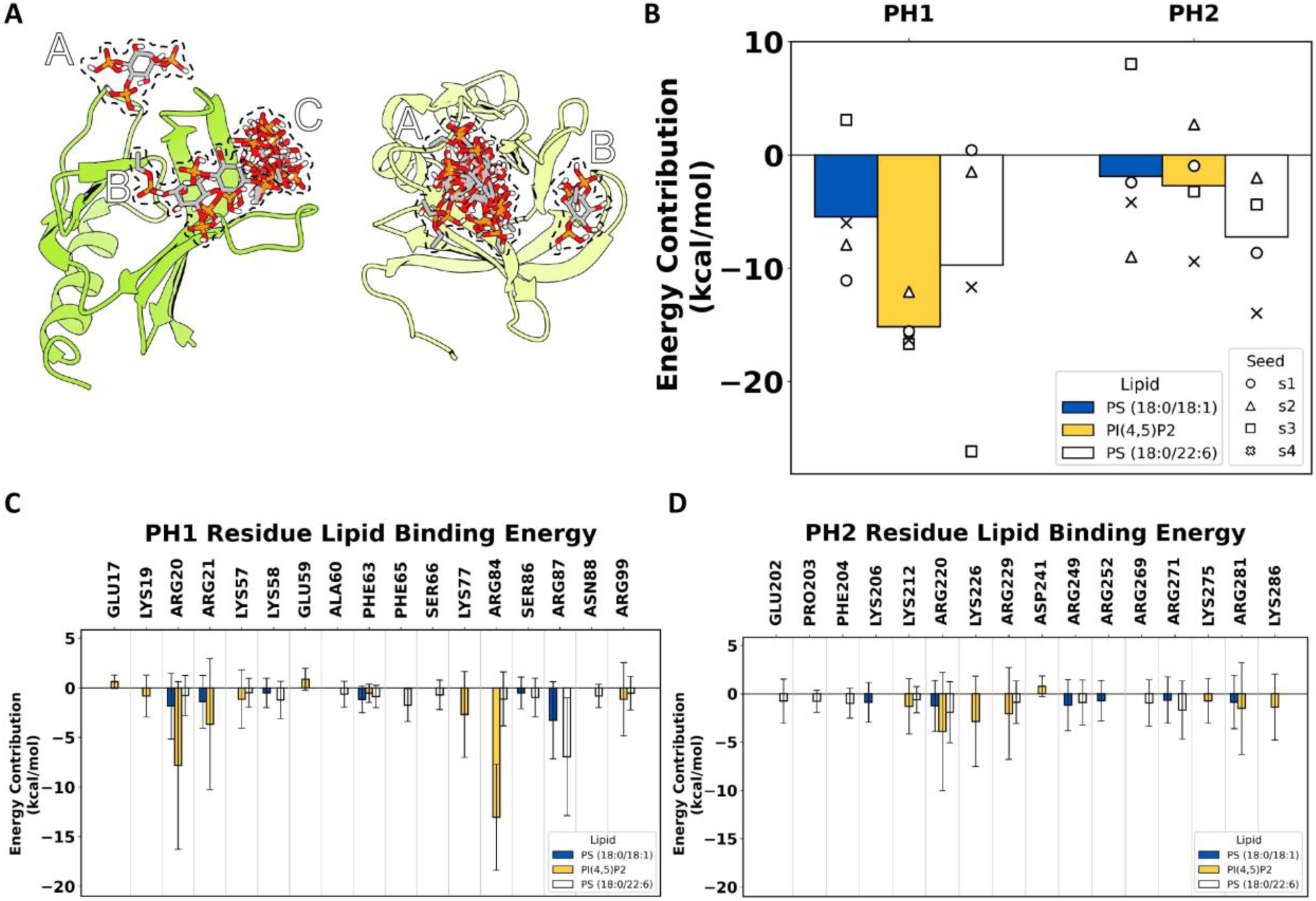
Both PH domains interact with lipids. **(A)** SwissDock results reveal three possible binding sites of PIP2 headgroup in PH1 (left) and two sites in PH2 (right). **(B)** MM/PBSA analysis of molecular dynamics simulations consisting of PH1 or PH2 and a PIP2-containing lipid bilayer. **(C)** PH1 residues involved in membrane binding. **(D)** PH2 residues involved in membrane binding.

**Supplementary Figure 4.**
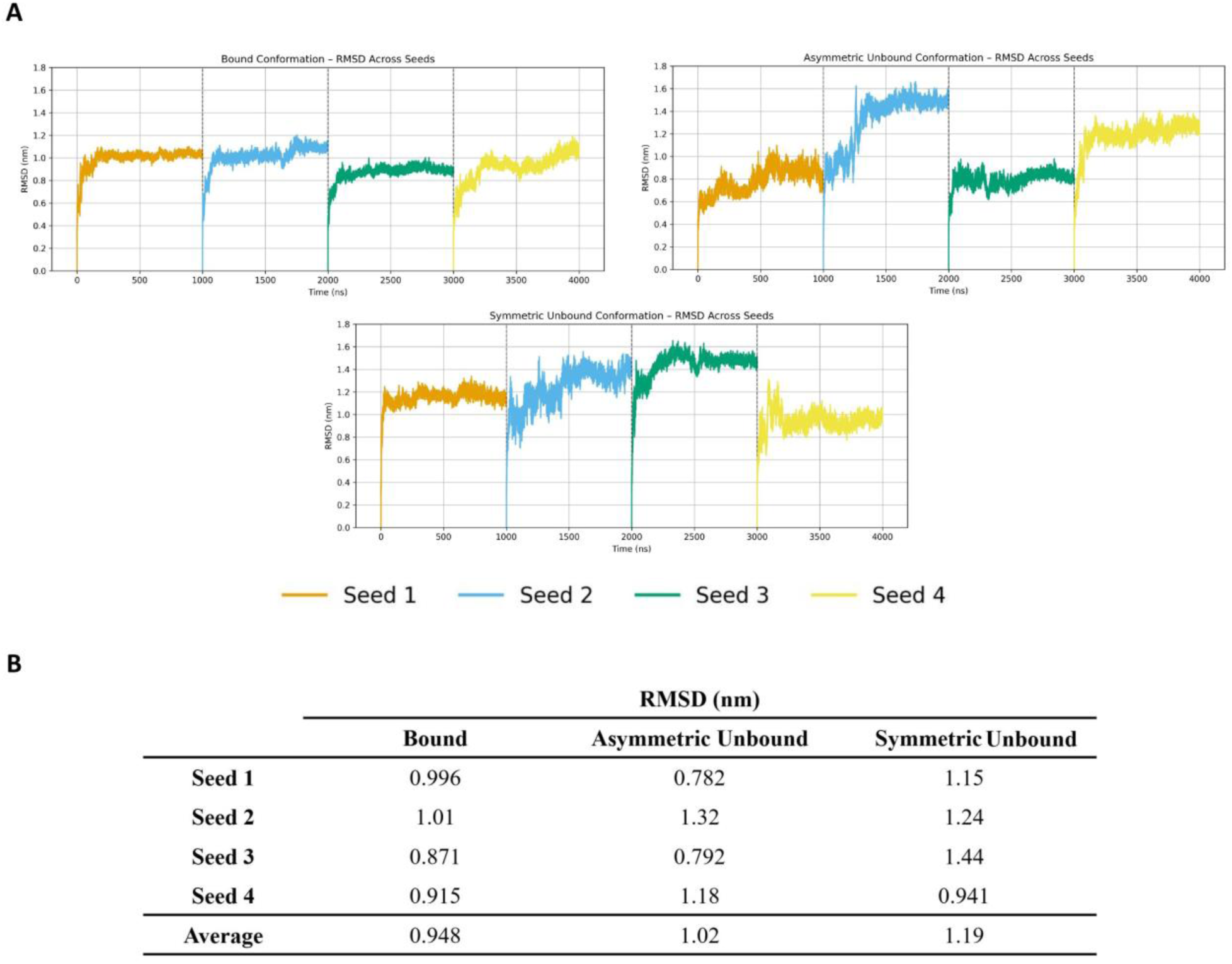
Molecular dynamics RMSD analysis. **(A)** RMSD of SNAP-47 across seeds for each conformation. Each seed is 1000 ns, and seeds are concatenated on each graph. **(B)** Average RMSD values across each seed for each conformation.

**Supplementary Figure 5.**
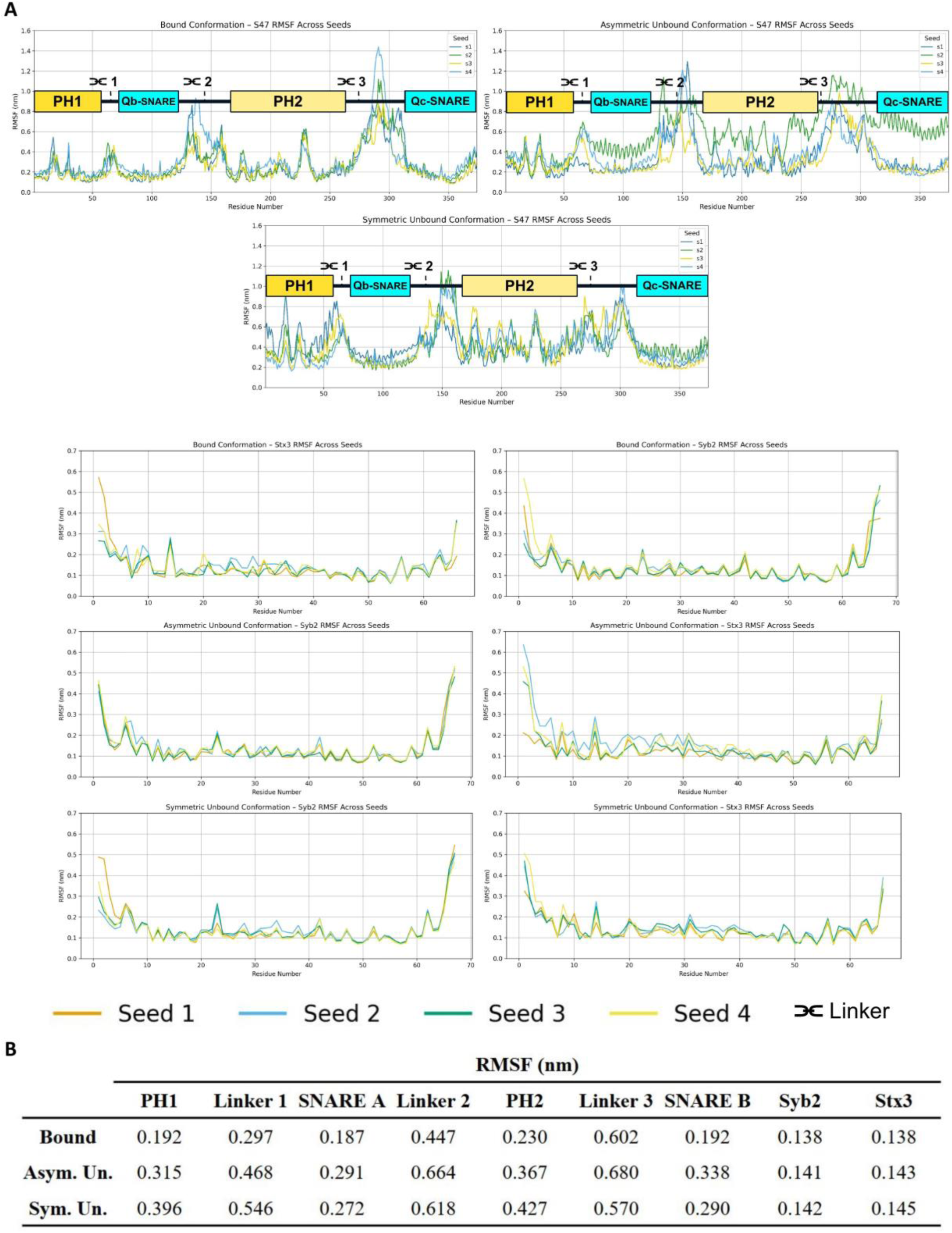
Molecular dynamics RMSF analysis. **(A)** RMSF of SNAP-47 (top), Stx-3 and Syb-2 (bottom) across seeds for each conformation. **(B)** Average RMSF values for each domain of SNAP-47 across each seed for each conformation.

**Supplementary Figure 6.**
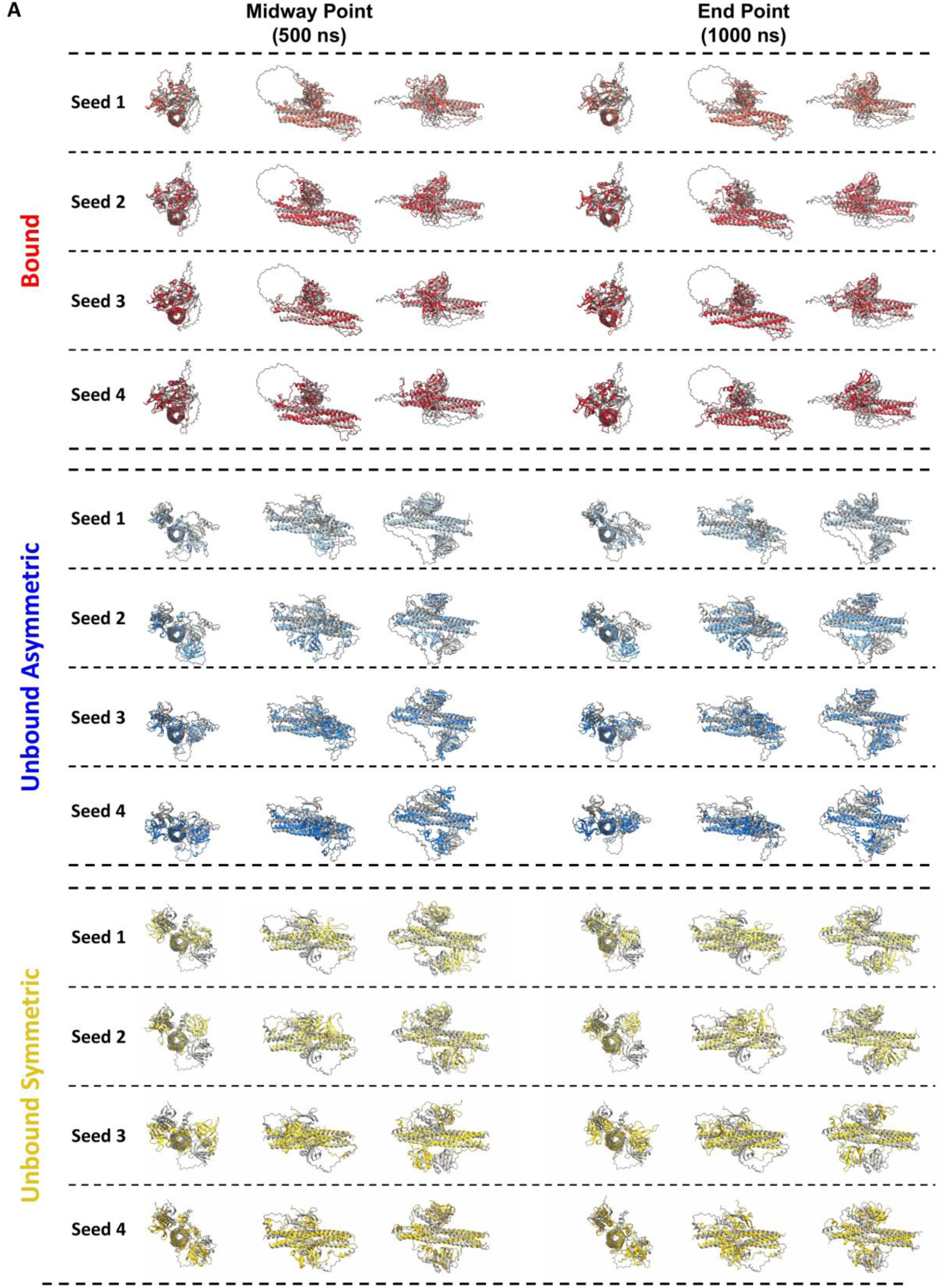
Structural overlays of molecular dynamics runs. **(A)** Overlay of the starting structure and the structure at the mid-point and endpoint of each trajectory for bound (top), unbound asymmetric (middle), and unbound symmetric (bottom) conformations.

**Supplementary Figure 7.**
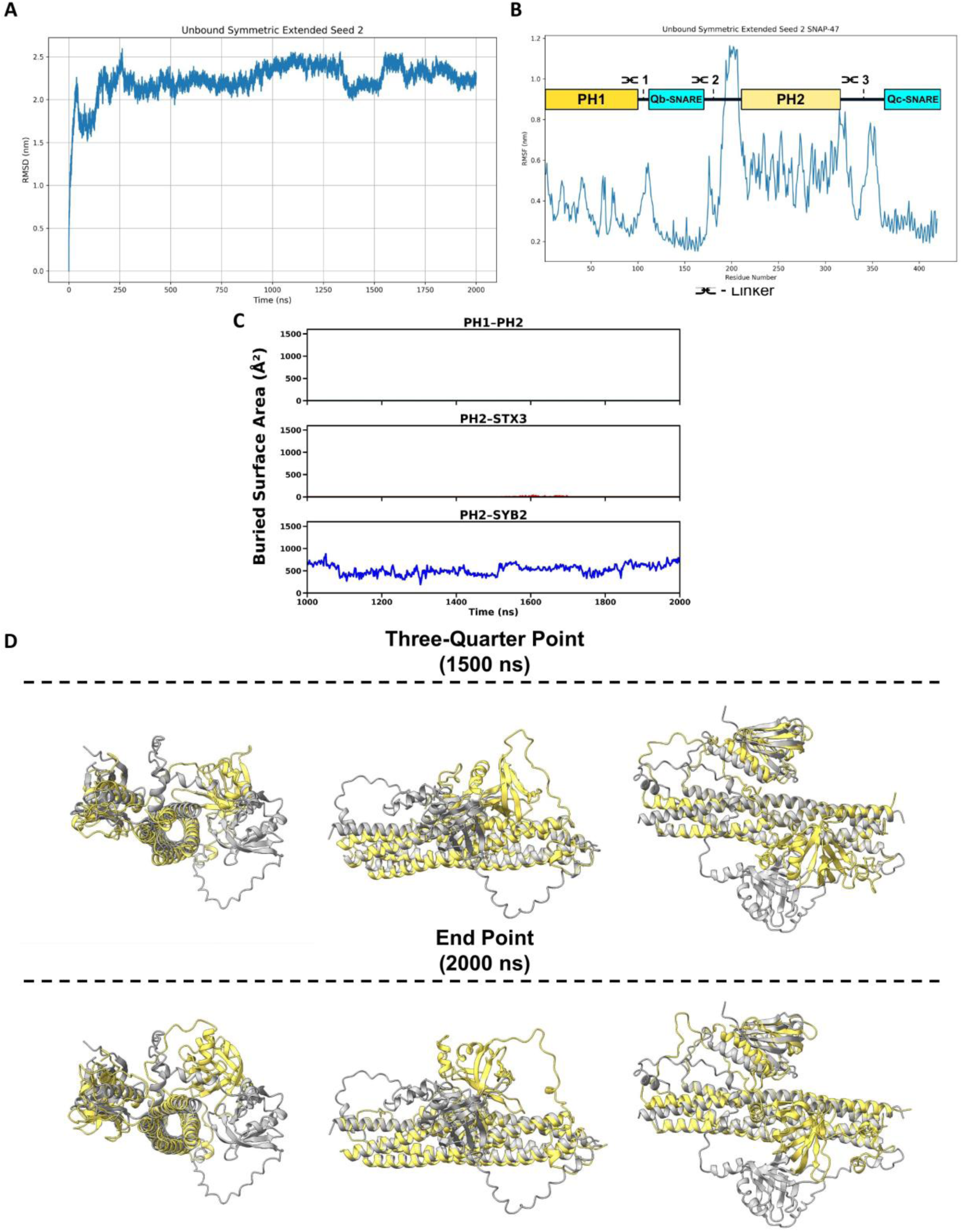
Extended unbound symmetric run. (A) RMSD and (B) RMSF results of the 1-µs extension of the original 1-µs unbound symmetric seed 2 simulation (2 µs total). **(C)** Buried surface area over time plot of the 1-µs extension for PH1—PH2, PH2—Stx-3, and PH2—Syb-2. **(D)** Overlay of structure at the midpoint and endpoint of 1 µs extension.

**Supplementary Figure 8.**
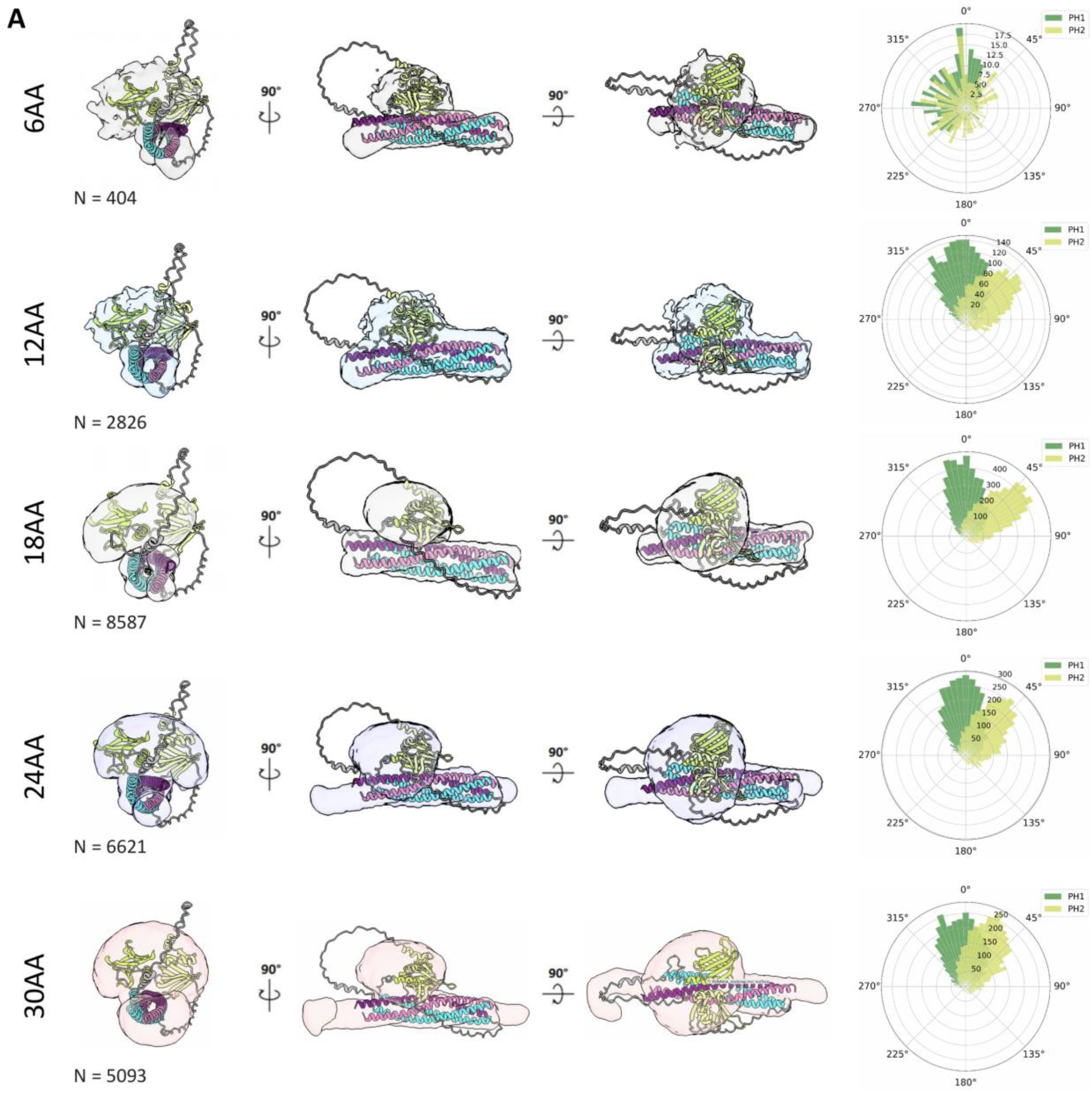
SNAP-47—Stx-3—Syb-2 SNARE complex BioEmu linker screen. **(A)** Surface maps corresponding to the primary cluster for each tested amino-acid GSG linker construct overlaid onto the AlphaFold-predicted bound conformation. Structures are shown in three views rotated 90° relative to one another, with a polar frequency plot on the right depicting the position of PH2 relative to PH1. N = number of structures that stably formed SNARE complexes out of the 10,000 predictions.

**Supplementary Figure 9.**
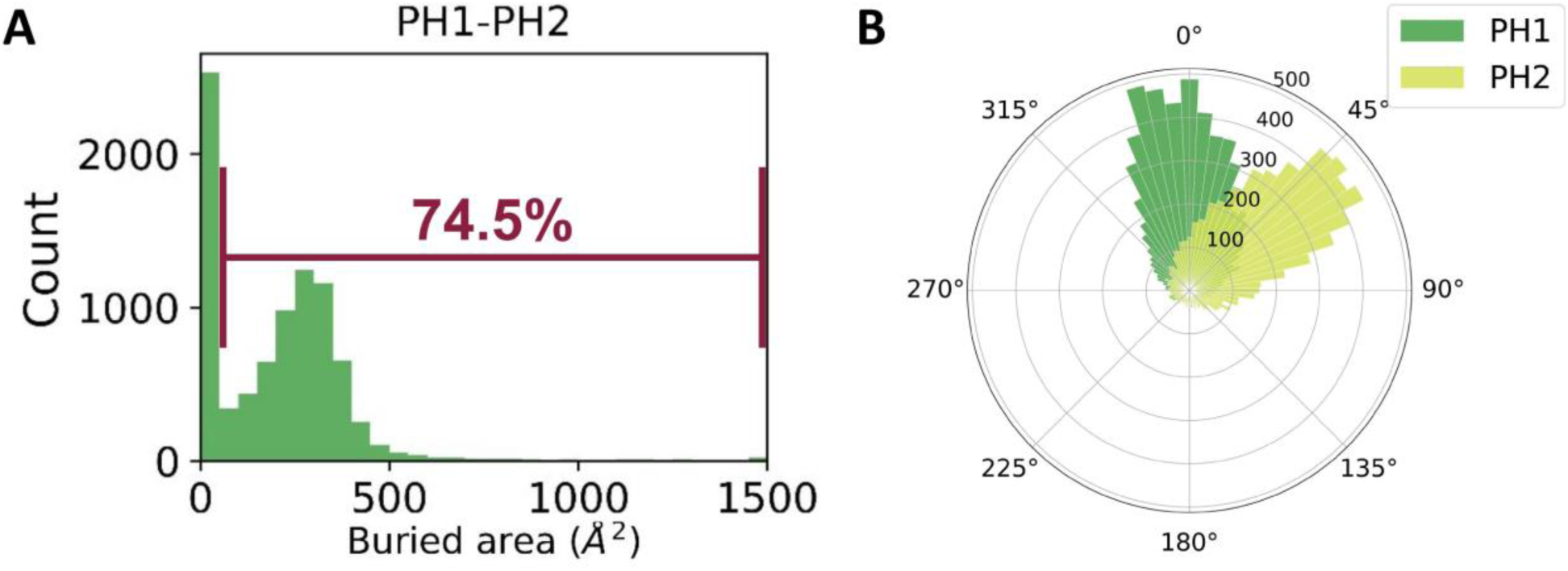
BioEmu shows formation of PH1—PH2 interface at angles consistent with bound conformation. **(A)** Histogram showing PH1—PH2 interface buried area for the 8,587 folded BioEmu predictions using the 18-amino-acid linker. **(B)** Polar graph showing PH domain angle relative to the SNARE coiled-coil structure for the 8,587 folded BioEmu predictions using the 18-amino-acid linker.

**Supplementary Figure 10.**
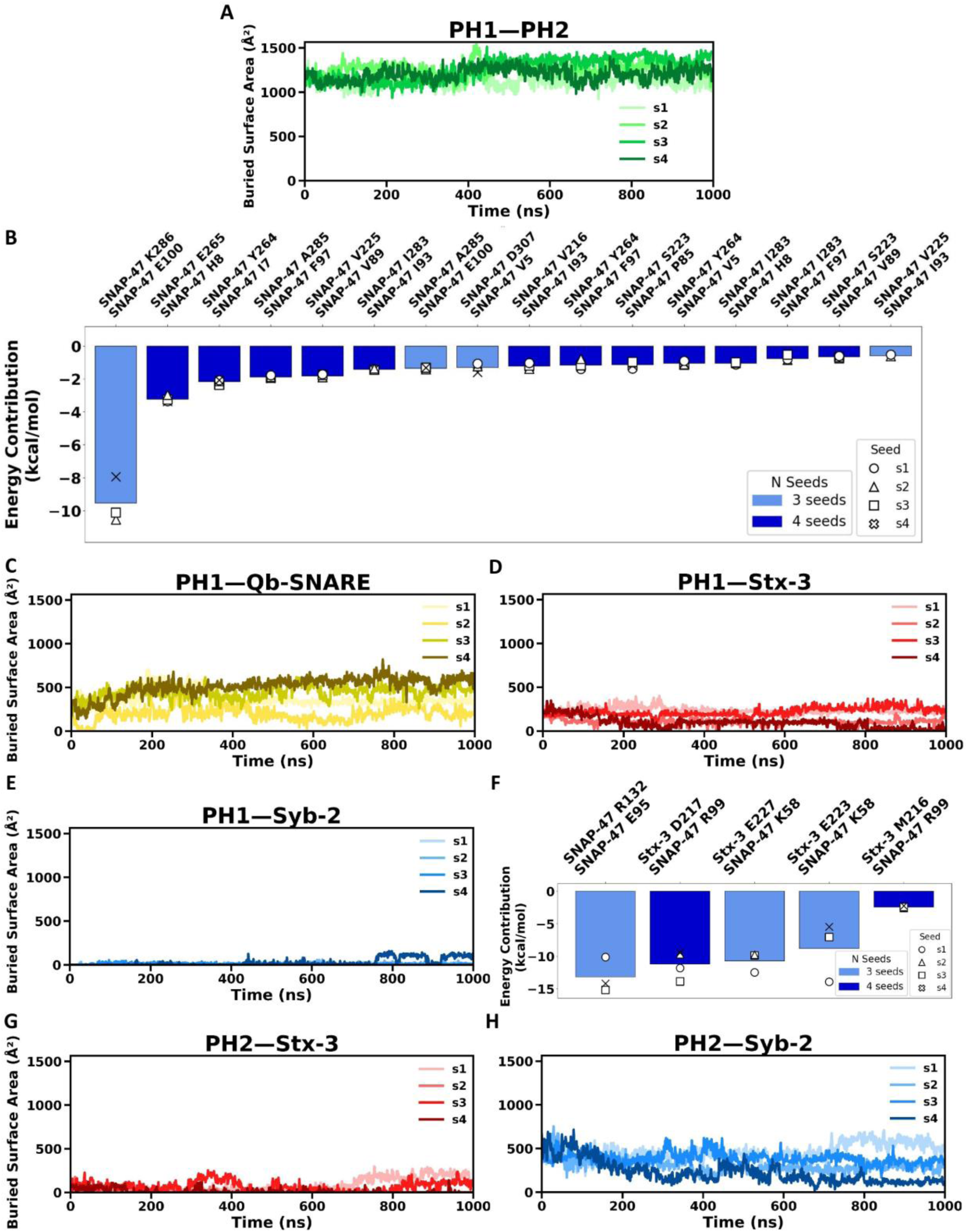
Buried surface area and MM/PBSA analyses of bound interfaces. **(A)** Buried Surface Area graph of PH1—PH2 across four seeds. **(B)** Top scoring residue pairs within PH1—PH2 present in three or more seeds as determined by MM/PBSA. **(C)** Buried Surface Area graph of PH1—Qb-SNARE across four seeds. **(D)** Buried Surface Area graph of PH1—Stx-3 across four seeds. **(E)** Buried Surface Area graph of PH1—Syb-2 across four seeds. **(F)** Top-scoring residue pairs within PH1—SNAREs present in three or more seeds as determined by MM/PBSA. **(G)** Buried Surface Area graph of PH2—Stx-3 across four seeds. **(H)** Buried Surface Area graph of PH2—Syb-2 across four seeds.

**Supplementary Figure 11.**
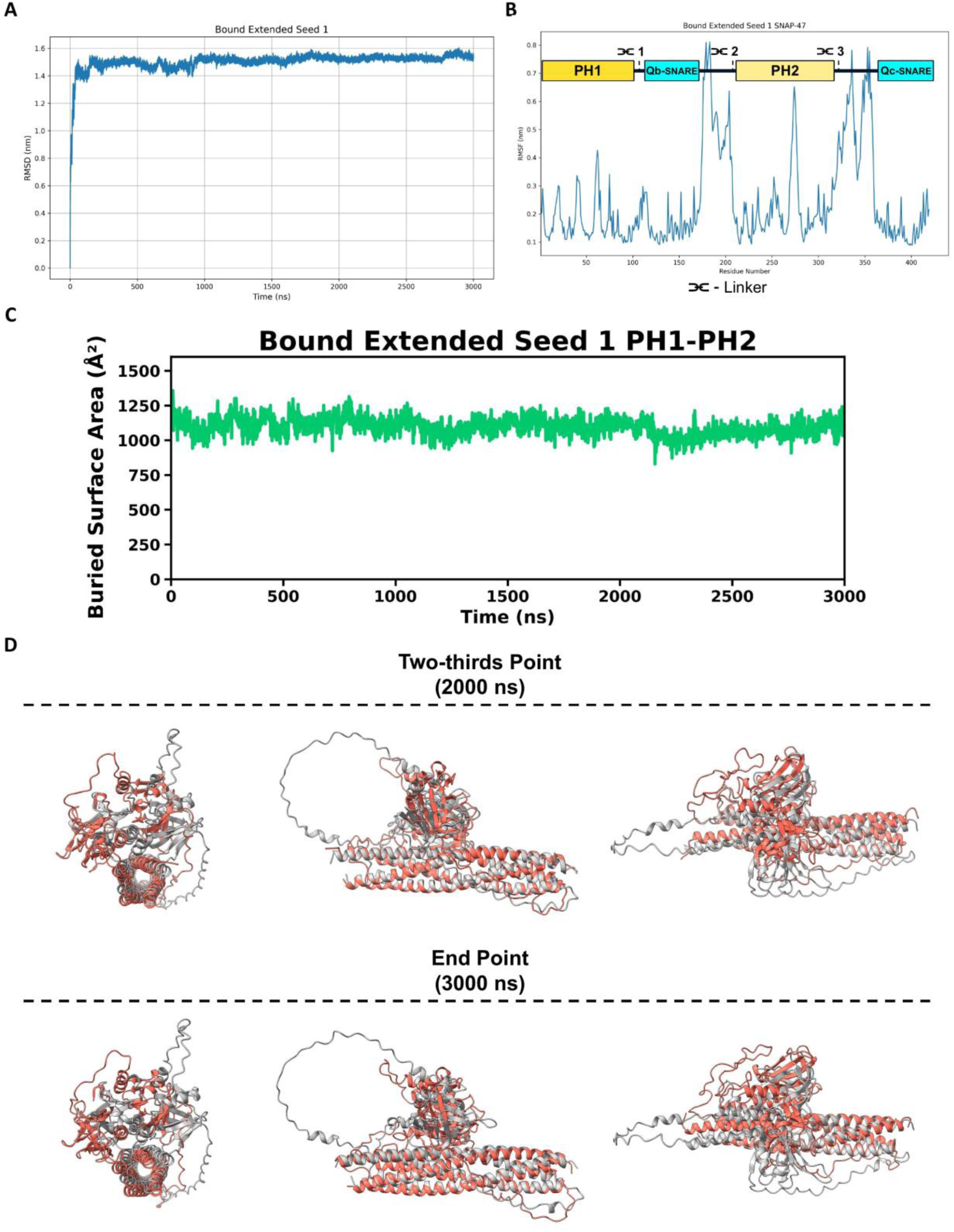
Extended bound MD simulation. (A) RMSD and (B) RMSF results of a 2-µs extension of the original 1-µs bound seed 1 simulation (3 µs total). **(C)** Buried surface area over time plot of the 2-µs extension for PH1—PH2. **(D)** Overlay of structure at the two-thirds and endpoint of the 2-µs extension.

**Supplementary Figure 12.**
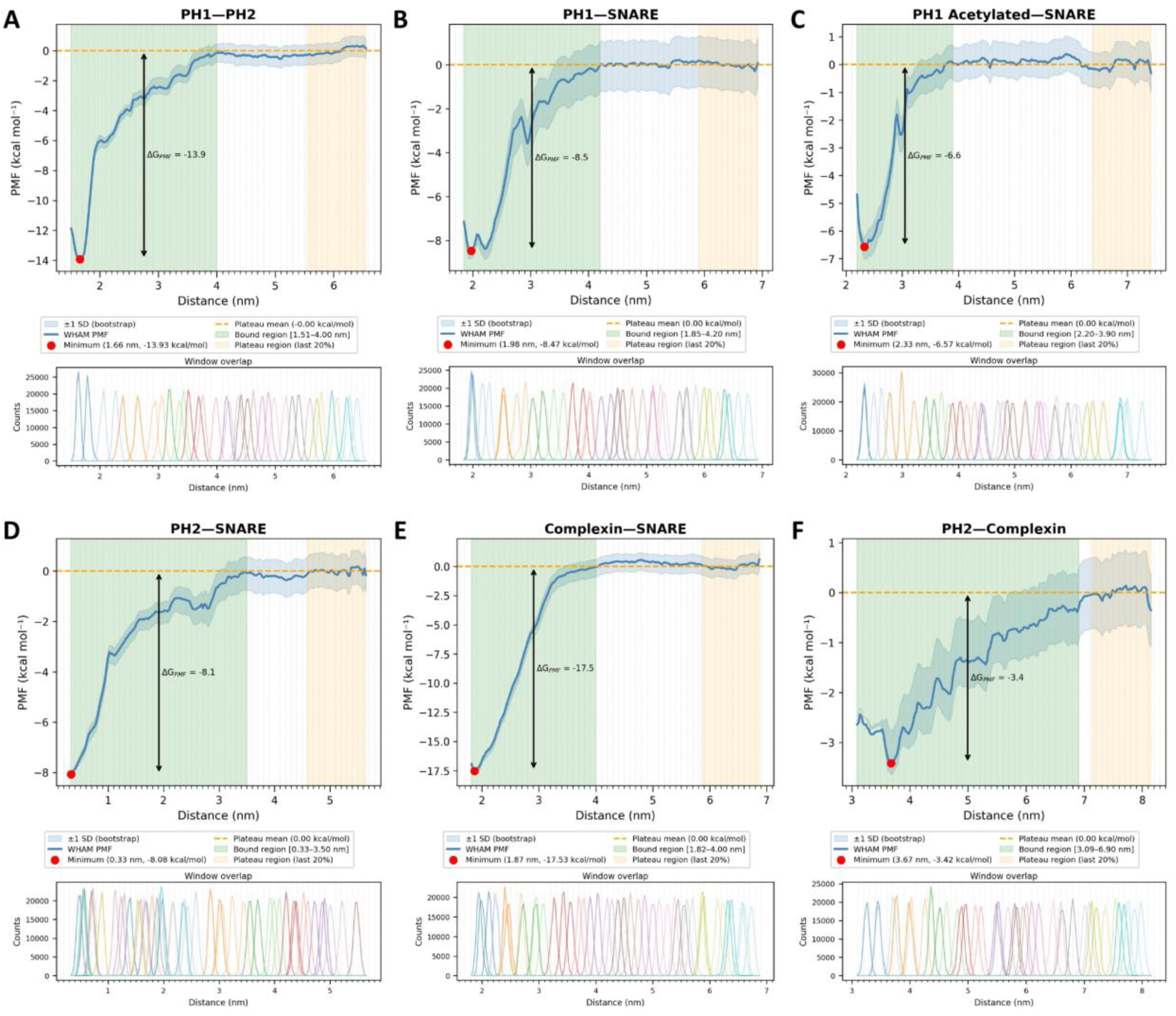
Umbrella sampling of top-scoring interfaces. Plots showing umbrella sampling PMF (top) and histogram coverage (bottom) over the reaction coordinate for **(A)** PH1—PH2, **(B)** PH1—SNARE, **(C)** PH1—SNARE acetylated, **(D)** PH2—SNARE, **(E)** Complexin—SNARE, and **(F)** PH2—complexin.

**Supplementary Figure 13.**
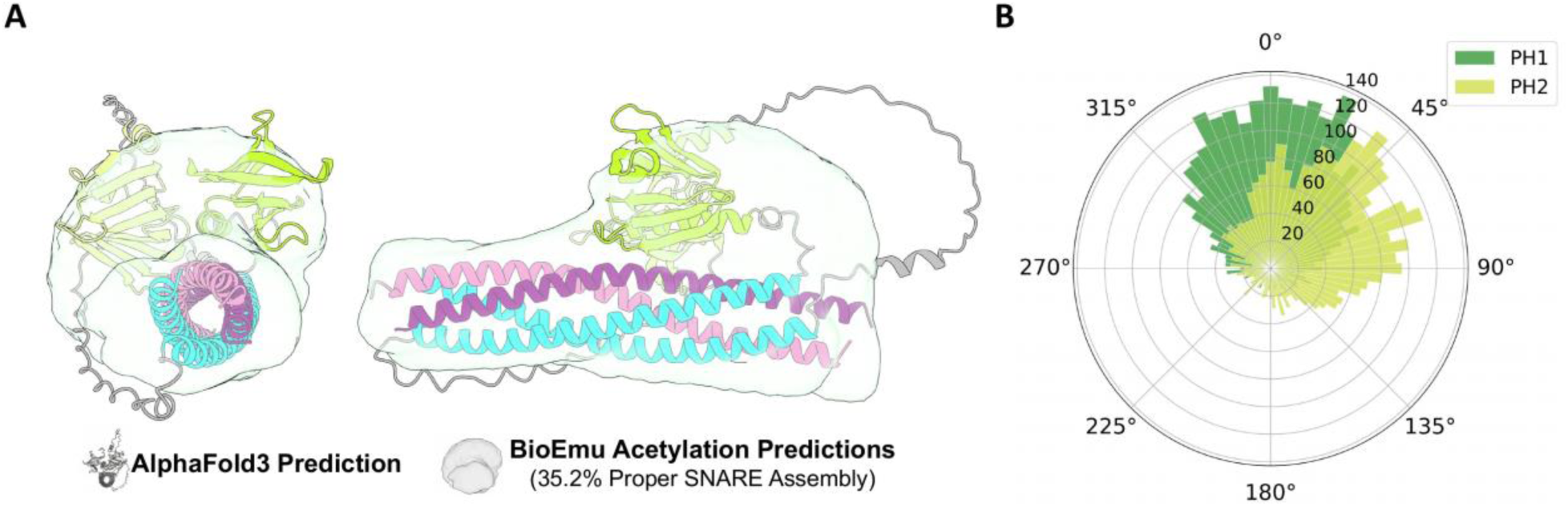
BioEmu acetylation. **(A)** BioEmu acetylation-mimic predictions shown as a gray voxel map of average occupancy. Voxel map is overlaid on the bound AlphaFold3 prediction. Structures are shown in two different views at a 90-degree rotation from each other. **(B)** Polar graph showing PH domain angle relative to SNARE coiled-coil structure.

**Supplementary Figure 14.**
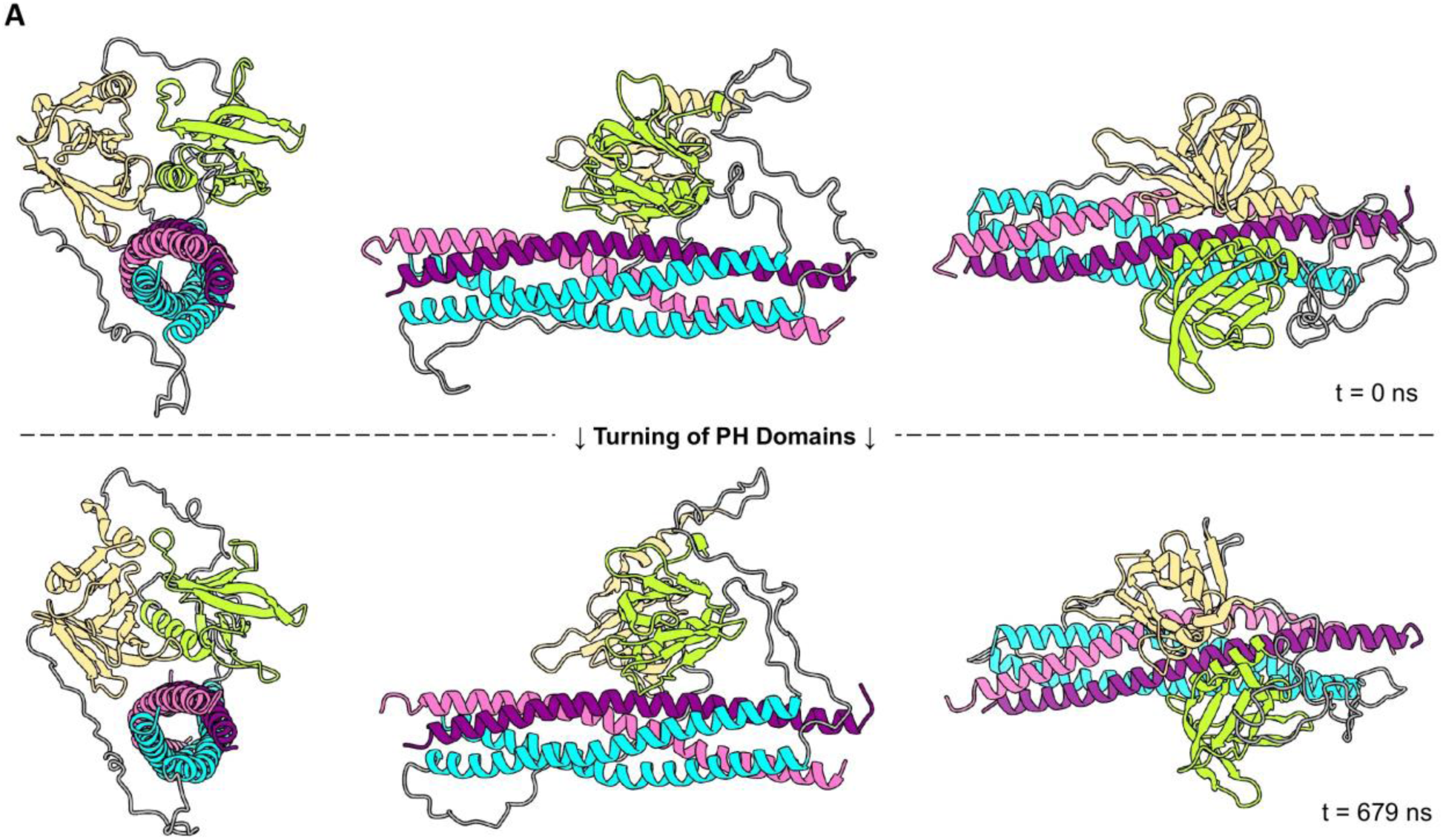
PH domain twist event observed in additional acetylation run. (A) Both PH1 and PH2 turn slightly from their starting position (top row) to a different angle within one of the acetylation seeds (bottom row).

**Supplementary Figure 15.**
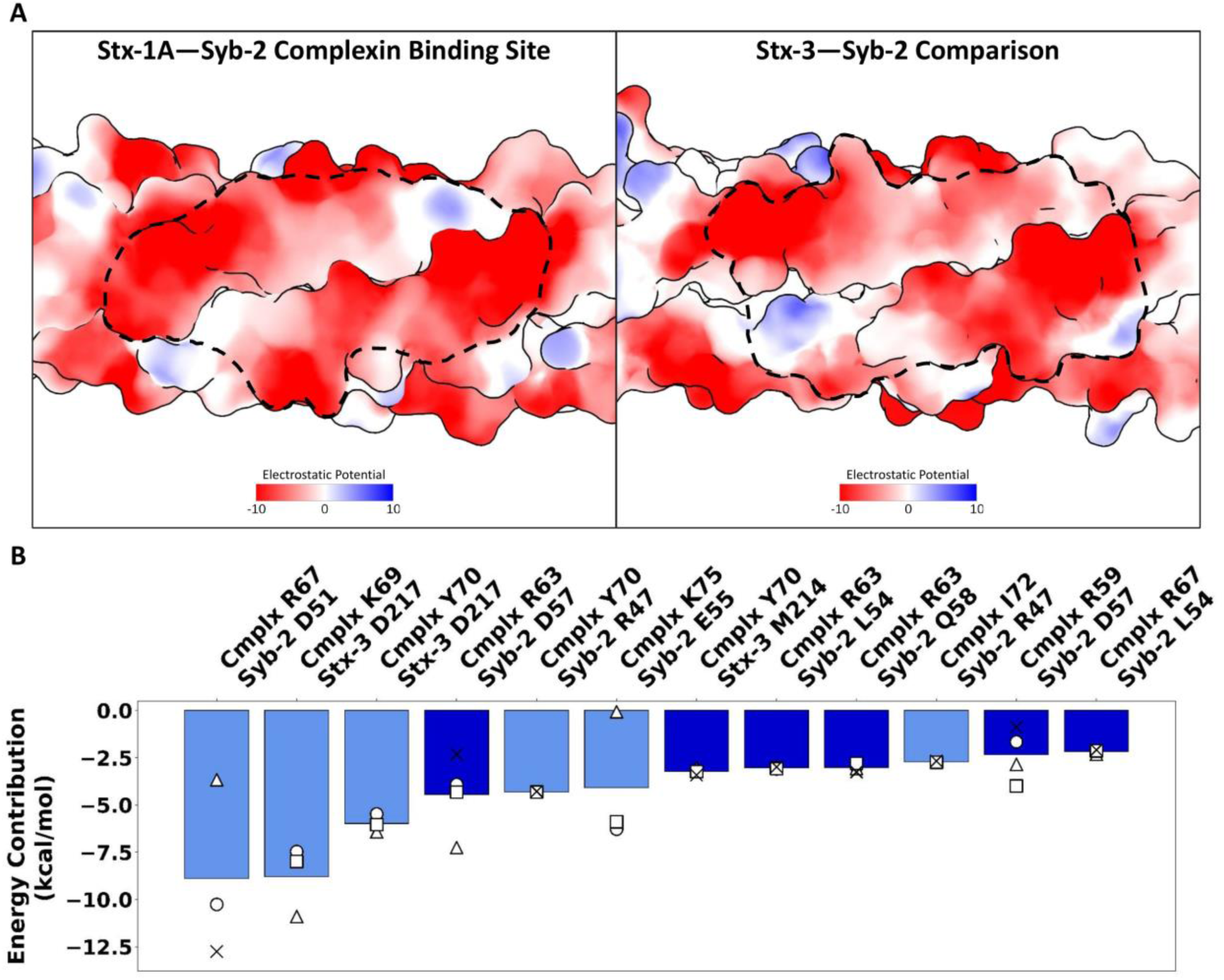
Residue pairs formed between complexin and SNAREs. **(A)** Electrostatic potential map showing Complexin’s binding site in the Stx-1a—Syb-2 groove (left) and the same groove within the Stx-3—Syb-2 system (right). **(B)** Top-scoring residue pairs within Complexin—SNAREs present in three or more seeds as determined by MM/PBSA.

**Supplementary Figure 16.**
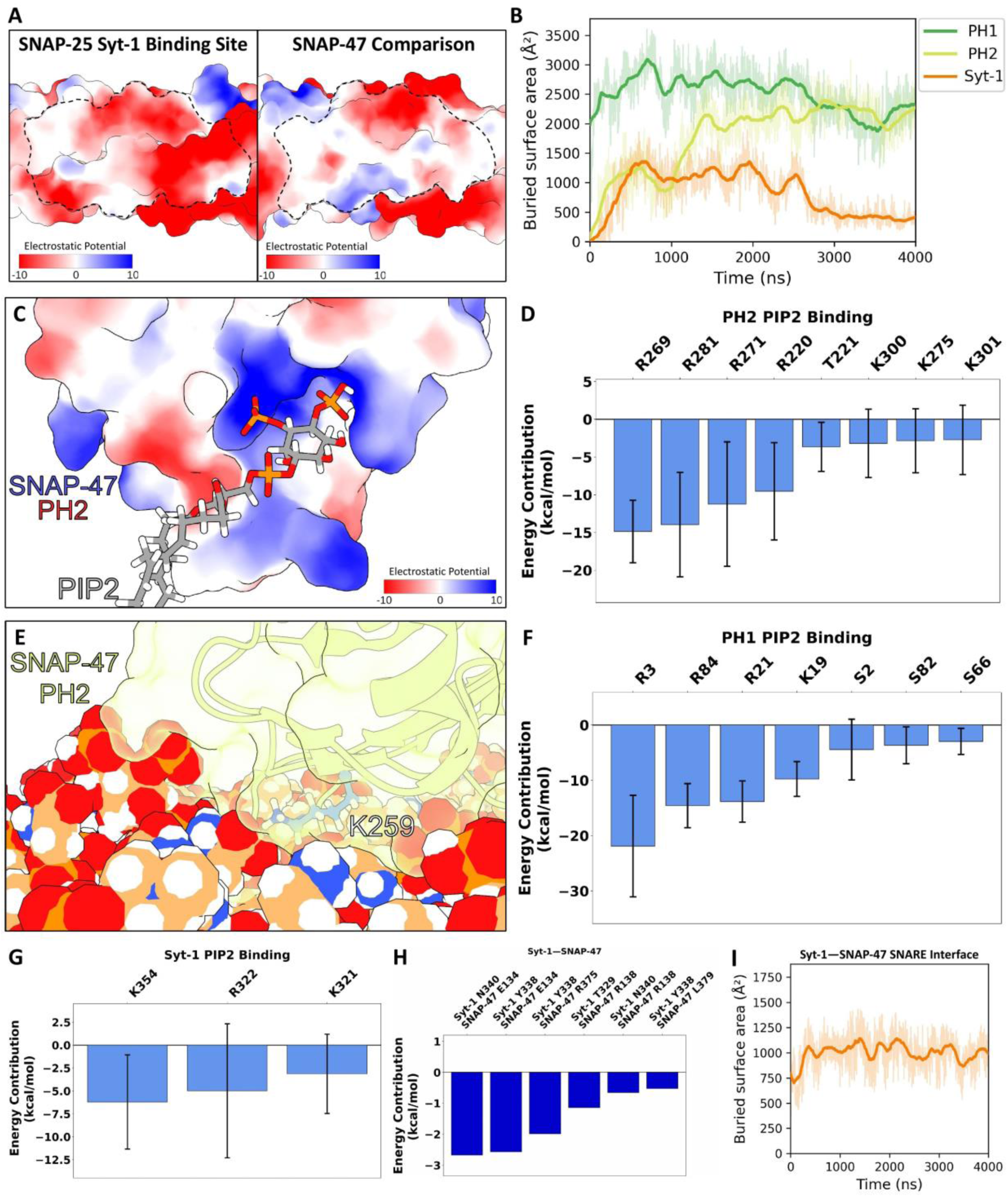
PH1, PH2, and Syt-1 C2B domains form interfaces with the plasma membrane in a 4-µs simulation. **(A)** Electrostatic potential map showing Syt-1’s binding site in the SNAP-25 SNARE groove (left) and the same groove in the SNAP-47 system (right). **(B)** Buried Surface Area plot showing PH1, PH2, and Syt-1 interacting with the membrane during the course of a 4-µs simulation. **(C)** Electrostatic map of the PH2 PIP2 binding pocket. **(D)** MM/PBSA results of PH2 and PIP2 binding. **(E)** Space-filling model showing that the additional SNAP-47 acetylation site K259 on PH2 is within the PH2—Membrane interface. **(F)** MM/PBSA results of PH1 and PIP2 binding. **(G)** MM/PBSA results of Syt-1 C2B and PIP2 binding. **(H)** MM/PBSA results of the Syt-1—SNAP-47 SNAREs interface. **(I)** Buried Surface Area for Syt-1—SNAP-47 SNARE.

**Supplementary Figure 17.**
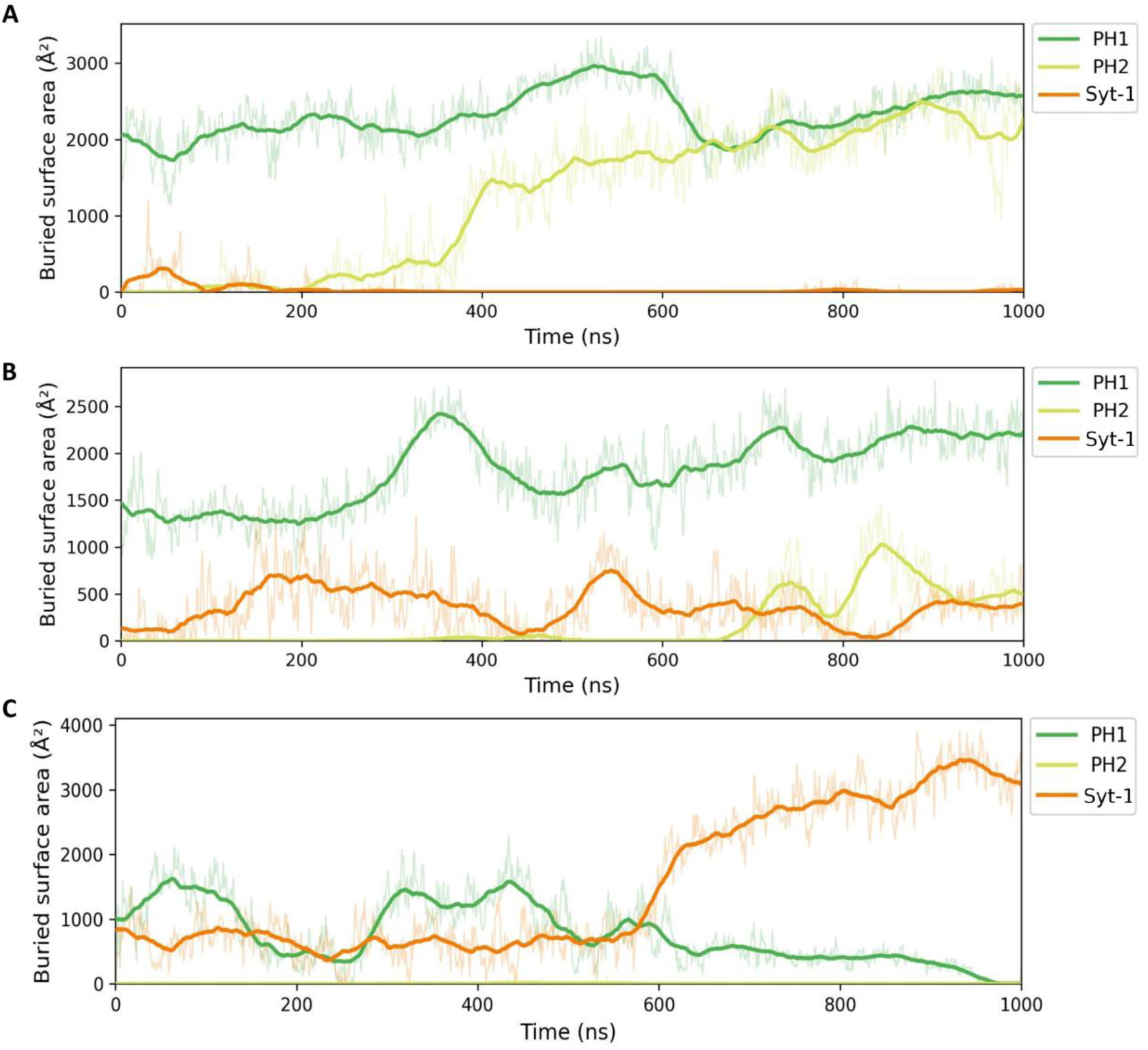
Buried surface area plots for additional SNARE complex simulations containing Syt-1 C2B and complexin. **(A)** Buried surface area plots for subsequent follow-up 1-µs simulations showing PH1 and PH2 only interacting with the membrane, **(B)** PH1, PH2, and Syt-1 interacting with the membrane, and **(C)** PH1 and Syt-1 only interacting with the membrane.

